# Single-fiber nucleosome density shapes the regulatory output of a mammalian chromatin remodeling enzyme

**DOI:** 10.1101/2021.12.10.472156

**Authors:** Nour J Abdulhay, Laura J Hsieh, Colin P McNally, Mythili Ketavarapu, Sivakanthan Kasinathan, Arjun S Nanda, Megan S Ostrowski, Ke Wu, Camille M Moore, Hani Goodarzi, Geeta J Narlikar, Vijay Ramani

## Abstract

ATP-dependent chromatin remodelers regulate the DNA accessibility required of virtually all nuclear processes. Biochemical studies have provided insight into remodeler action at the nucleosome level, but how these findings translate to activity on chromatin fibers *in vitro* and *in vivo* remains poorly understood. Here, we present a massively multiplex single-molecule platform allowing high-resolution mapping of nucleosomes on fibers assembled on mammalian genomic sequences. We apply this method to distinguish between competing models for chromatin remodeling by the essential ISWI ATPase SNF2h: linker-length-dependent dynamic positioning versus fixed-linker-length static clamping. Our single-fiber data demonstrate that SNF2h operates as a density-dependent, length-sensing chromatin remodeler whose ability to decrease or increase DNA accessibility depends on single-fiber nucleosome density. *In vivo*, this activity manifests as different regulatory modes across epigenomic domains: at canonically-defined heterochromatin, SNF2h generates evenly-spaced nucleosome arrays of multiple nucleosome repeat lengths; at SNF2h-dependent accessible sites, SNF2h slides nucleosomes to increase accessibility of motifs for the essential transcription factor CTCF. Overall, our generalizable approach provides molecularly-precise views of the processes that shape nuclear physiology. Concurrently, our data illustrate how a mammalian chromatin remodeling enzyme can effectively sense nucleosome density to induce diametrically-opposed regulatory effects within the nucleus.

## INTRODUCTION

Nucleosomes regulate the DNA transactions essential to life. Continuous, yet cell-type specific, rearrangement of nucleosomes is carried out by many *trans*-acting factors, including ATP-dependent chromatin remodeling complexes (*i.e*. ‘chromatin remodelers’). Chromatin remodelers fall under four evolutionarily-conserved families of DNA translocase—SWI/SNF, ISWI, INO80, and CHD—each of which harness the energy of ATP to slide, evict, load, or transfer core histones (Narlikar et al., 2013). Remodelers, which generally operate in the context of multimeric protein complexes, are often developmentally essential and highly dosage-sensitive, and genetic mutations in catalytic and non-catalytic remodeler subunits have been implicated in a variety of diseases, including cancer (Kadoch, 2019; Kadoch et al., 2013; Lek et al., 2016).

Our current understanding of how sequence-non-specific chromatin remodeling complexes act on chromatin derives from a combination of biochemical studies on mononucleosomes, and bulk nucleosome footprinting studies on oligonucleosomes (*i.e*. ‘chromatin fibers’). However, nucleosome positions, and the average nucleosome spacing on individual chromatin fibers, can vary substantially across a population of even identical DNA templates. As such, understanding how remodelers operate in the context of individual chromatin fibers is critical to evaluating their biological role regulating DNA accessibility.

Unfortunately, existing bulk footprinting methods cannot capture the connectivity and heterogeneity of nucleosome positions within a chromatin fiber. To bridge this major experimental gap, we develop and apply a high-resolution single-fiber approach, drawing inspiration from a series of seminal studies wherein remodeling activities were surveyed against *in vitro* chromatinized genomic DNA templates using high-throughput genomic footprinting, an approach referred to as ‘genomic reconstitution’ (Kaplan et al., 2009; Krietenstein et al., 2016; Zhang et al., 2011). Specifically, we couple genomic reconstitution with our previously published, <u>s</u>ingle-molecule <u>a</u>denine <u>m</u>ethylated <u>o</u>ligonucleosome <u>s</u>equencing <u>a</u>ssay (SAMOSA) (Abdulhay et al., 2020), and use this new approach to study ISWI-mediated nucleosome remodeling at single chromatin fiber resolution (Tsukiyama et al., 1995).

Mammalian ISWI complexes catalyze nucleosome sliding via the ATPase motors sucrose nonfermenting 2 homologue / homologue-like (SNF2h / SNF2l). These ATPases partner with non-catalytic subunits to promote many essential nuclear functions, including DNA replication, repair, transcriptional activation, and repression (Erdel and Rippe, 2011). While ISWI complexes are known to organize nucleosomes into equally-spaced arrays, the manner by which ISWI complexes equalize spacing remains debated. There are currently two distinct models for how ISWI remodelers define internucleosomal distances (IDs) on chromatin fibers (see **Figure 3A**). First, a ‘clamping’ (also known as ‘ruler’) model proposes that SNF2h slides nucleosomes to create fixed IDs as if via a ‘clamp.’ The HAND-SAND-SLIDE (HSS) domain of SNF2h, has been proposed to enable such clamping by binding a defined length of linker DNA. In this model IDs are largely independent of the underlying nucleosome density (# of nucleosomes per unit length DNA) of the chromatin fiber (Krietenstein et al., 2016; Lieleg et al., 2015; Oberbeckmann et al., 2021). Second, a linker-length-dependent model (*i.e*. a ‘length-sensing’ model) proposes that SNF2h instead uses the HSS to sense the length of extranucleosomal flanking DNA, and slides nucleosomes faster in the direction of longer flanking DNA. In this model, ISWI enzymes sense differences in linker lengths up to the maximal linker length bound by the HSS (Blosser et al., 2009; Deindl et al., 2013; Leonard and Narlikar, 2015; Yang et al., 2006). The length-sensing model predicts that steady-state linker lengths generated by SNF2h will depend on pre-existing chromatin density, and that at sufficiently low densities, SNF2h will isotropically translocate nucleosomes along template DNA. Distinguishing between these models may be essential to understanding how ISWI promotes silencing at specific sites, while exposing others to reveal *cis*-regulatory motifs for essential transcription factors (TFs) like CTCF (Barisic et al., 2019; Wiechens et al., 2016).

Studies performed to-date harbor unique limitations that confound resolution of either model. Bulk and single-molecule studies, for instance, have been performed in the context of mononucleosomes (Blosser et al., 2009; Racki et al., 2009; Yang et al., 2006) while *in vitro* activity measurements on chromatin fibers have relied on bulk nuclease digestion (Krietenstein et al., 2016; Lieleg et al., 2015; Oberbeckmann et al., 2021; Zhang et al., 2011). Delineation between these models is further complicated by the facts that i.) equally-spaced nucleosome arrays can randomly emerge downstream of a barrier without invoking nucleosome remodeling (*i.e*. ‘statistical’ positioning) (Kornberg and Stryer, 1988), and ii.) primary sequence can influence initial nucleosome positions (Lowary and Widom, 1998).

The single-fiber method we present here is uniquely suited to overcoming these limitations. Using our approach, we find that SNF2h activity is consistent with the length-sensing model, as average SNF2h-catalyzed nucleosome spacing on single fibers decreases as underlying nucleosome density increases. We explore the physiological significance of this model using binding motifs for the TF CTCF, demonstrating that the initial nucleosome-location and density ‘states’ of a fiber ultimately dictate the fold-change in relative accessibility of sites following remodeling. Finally, to understand implications of this length-sensing model *in vivo*, we footprint individual chromatin fibers in living murine embryonic stem cells (mESCs) devoid of SNF2h. Our results demonstrate that at heterochromatic sites known to be targeted by ISWI complexes, SNF2h acts to generate a population of evenly-spaced nucleosome arrays with short, but variant, nucleosome repeat lengths; conversely, at promoters and Ctcf binding sites, ISWI complexes increase DNA accessibility by sliding nucleosomes to create irregular fibers. Our single-fiber experiments rule out the clamping model for SNF2h activity *in vitro* and *in vivo*, suggesting that ISWI remodelers sense flanking DNA length and nucleosome density to enact multiple regulatory functions. Taken as a whole, our study illustrates how nucleosome sliding in the context of varying nucleosome density can program DNA accessibility, and offers a new paradigm for studying single-fiber chromatin remodeling.

## RESULTS

### Single-molecule footprinting of intact chromatin fibers reconstituted on genomic sequences

Prior biochemical studies on chromatin fibers have used arrays composed of a nucleosome positioning sequence such as Widom 601 (Dechassa et al., 2010; Mivelaz et al., 2020), or fibers reconstituted from yeast genomic DNA (Kaplan et al., 2009; Krietenstein et al., 2016; Zhang et al., 2011). We previously demonstrated that our SAMOSA protocol could accurately resolve single-fiber nucleosome footprints on 601-based chromatin fibers (Abdulhay et al., 2020). To enable study of more native-like fibers, we extended the SAMOSA approach to footprint individual fibers reconstituted on mammalian genomic sequences through salt gradient dialysis (SGD). To this end, we devised a general workflow we term SAMOSA-ChAAT (<u>Ch</u>romatin <u>A</u>ccessibility of <u>A</u>ssembled <u>T</u>emplates; **Figure 1A**), in which chromatin fibers with desired biochemical properties (*e.g*. nucleosome density) are assembled from genomic templates, subjected to the SAMOSA m^6^dA footprinting protocol, and sequenced on the PacBio Sequel II to natively detect m^6^dA modifications reflective of accessible DNA bases.

**Figure 1:**
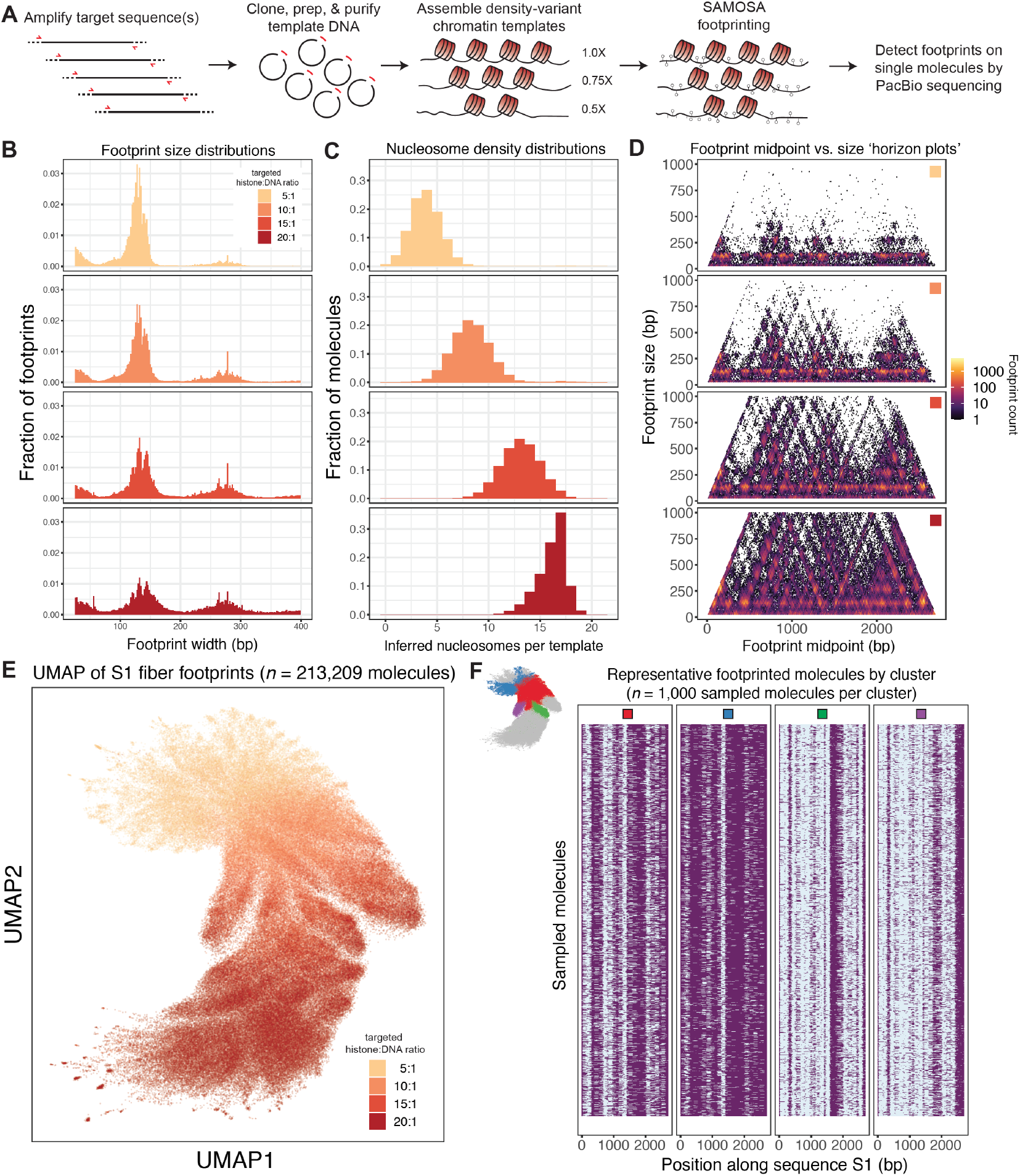
SAMOSA-ChAAT enables massively multiplex dissection of single-fiber nucleosome positioning on *in vitro* reconstituted genomic chromatin fibers. **A.)** Schematic overview of the SAMOSA-ChAAT protocol, wherein genomic sequences are cloned, purified, and assembled into chromatin fibers with desired biochemical properties (*e.g.* nucleosome density) through salt gradient dialysis. Fibers are then footprinted with a nonspecific adenine methyltransferase and sequenced on the PacBio platform to assess single-molecule nucleosome positioning. **B.)** A custom analytical pipeline enables detection of methyltransferase footprints on sequenced fibers. Footprint sizes from SAMOSA-ChAAT experiments carried out at varying nucleosome densities follow closely with expected nucleosome sizes, plus expected ‘breathing’ of DNA around the histone octamer, with the extent of breathing decreasing as nucleosome density increases. **C.)** SAMOSA-ChAAT data enables direct estimation of the absolute number of nucleosomes per footprinted fiber, which track well with expected nucleosome densities based on targeted octamer: DNA ratios during salt gradient dialysis. **D.)** Footprint length vs. midpoint ‘horizon’ plots for footprinted fibers. Average nucleosome positions display sequence dependencies. **E.)** UMAP dimensionality reduction of fiber accessibility data. UMAP patterns recapitulate known differences in nucleosome density in footprinted fibers. **F.)** Visualization of a subset of sampled molecules following Leiden clustering of single molecule data. Individual Leiden clusters (cluster positions inset) capture mutually exclusive nucleosome positions consequent of chromatin fiber assembly.

As proof-of-concept for this approach, we cloned two ~3 kilobase sequences from the *M. musculus* genome (hereafter, sequences ‘S1’ and ‘S2’), carried out the SAMOSA-ChAAT workflow across four specified histone octamer:DNA ratios, and sequenced resulting molecules and controls to high depth (samples and sequencing depths summarized in **Supplementary Table 1**). We then analyzed the resulting sequencing data using a custom pipeline that combines a neural network with a hidden markov model (NN-HMM; **Methods**) to detect stretches of unmethylated DNA directly from PacBio data, while accounting for primary sequence biases for the EcoGII m^6^dAase and PacBio sequencing polymerase (**Supplementary Figure 1**). Consistent with the assembly of histones into nucleosome core particles with varying degrees of ‘breathability,’ we were able to call stretches of unmethylated DNA on sequenced molecules (*i.e*. ‘footprints’) ranging from ~120 – 160 nucleotides (nts) in size (**Figure 1B**), in addition to short (< 30 nt) footprints suggestive of nonspecific histone-DNA interactions (*e.g*. H2A/H2B-DNA). Footprint sizes increased along with chromatin density, suggesting i.) that higher nucleosome densities inhibit DNA breathing along assembled nucleosome core particles (footprint sizes for S1 predicted mononucleosomes: 130 ± 10.4 bp to 141 ± 19.3 bp for 5:1 to 20:1 fibers; data reported as median ± median absolute deviation), and ii.) that higher nucleosome densities promote formation of closely spaced di- and tri-nucleosome structures. Our data also enable estimates of the number of nucleosomes per individual template (*i.e*. ‘nucleosome density’); accordingly, inferred nucleosome counts on single molecules closely matched targeted assembly extents (**Figure 1C**; values reported as mean ± standard deviation in **Supplementary Table 2**).

Nucleosome assembly can be influenced by the underlying shape and rigidity of template DNA, which varies strongly as a function of DNA sequence (Rohs et al., 2009). To ascertain patterns of favored nucleosome positioning in bulk, we generated footprint length versus footprint midpoint ‘horizon plots’ (analogous to fragment length vs midpoint ‘V-plots’ (Henikoff et al., 2011)) for each assembly condition and sequence (**Figure 1D**). Our approach allows for explicit mapping and classification of footprints of all sizes as a function of target sequence, clearly revealing both sequence-directed nucleosome positioning, and regions that favor formation of closely-packed primary structures (*e.g*. dinucleosomes with virtually no intervening linker DNA).

To move beyond these bulk averages, we next explored our data at single-molecule resolution (**Figure 1E,F**) using UMAP dimensionality reduction (McInnes et al., 2018) and Leiden community detection (Traag et al., 2019). We found i.) that UMAP projections natively capture the biochemical parameter of chromatin density (**Figure 1E**), and ii.) that unbiased clustering enables detection of mutually exclusive nucleosome positions for molecules from identical SGD preparations (see purple and green clusters in **Figure 1F**). Importantly, our data satisfy a wide set of controls. First, our footprint-size analyses, nucleosome-density measurements, horizon plot visualizations, UMAP reductions, and cluster profiles were all consistent for the completely different sequence S2 (**Supplementary Figure 2**). Second, our analytical pipeline accurately detected expected footprint sizes and positions from Widom 601 chromatin fibers with known dyad positions (**Supplementary Figure 3A-B**), albeit with a longer mononucleosome footprint size consistent with less DNA breathing on Widom 601 nucleosomes (Lowary and Widom, 1998). Finally, our nucleosome occupancy measurements were highly quantitatively reproducible across replicates (**Supplementary Figure 3C-F**). Together, these data demonstrate the sensitivity, reproducibility, and generalizability of the SAMOSA-ChAAT approach.

### SAMOSA-ChAAT captures single-fiber chromatin remodeling reaction outcomes

We next used SAMOSA-ChAAT to study ATP-dependent chromatin remodeling by purified SNF2h at single-fiber resolution. Our multiplexing strategy allowed us to study multiple reaction conditions in parallel, including a pre-catalytic state (SNF2h(-)ATP), a remodeled steady state (SNF2h(+)ATP), and an uncatalyzed state where ADP is added instead of ATP (SNF2h(+)ADP), which we collectively compared against unremodeled chromatin fibers (*i.e*. ‘native’ fibers). The endpoint of 15 minutes was chosen to enable reaction completion (> 3 half-times) based on previously measured rate-constants (Yang et al., 2006). The vast majority of data we collected were under single-turnover and saturating reaction conditions ([SNF2h] ≫ [template]), though we also performed a small subset of control experiments under multiple turnover conditions where [SNF2h] < [template]. Including all replicate and control experiments, and after filtering out molecules that failed quality-control, we analyzed 3.17E6 footprinted molecules across multiple sequencing runs, amounting to a single-molecule fold-coverage of 1.72E6X and 1.45E6X for templates S1 and S2, respectively (**Supplementary Table 2**). As with our initial SGD preps, bulk repositioning of nucleosomes was highly quantitatively reproducible by our footprinting assay (**Supplementary Figure 4A-H**).

We focused on exploring the effects of remodeling on ‘10:1’ density S1 and S2 chromatin fibers. First, we visualized the bulk consequences of remodeling fibers through horizon plots (**Figure 2A-B; Supplementary Figure 4I-J**). Prior studies with mononucleosomes have shown that ATP-dependent chromatin remodelers move nucleosomes off from strong and weak positioning sequences with similar rates (Partensky and Narlikar, 2009). We found that SNF2h remodeling also decreases sequence-dependent nucleosome positioning on fibers— nucleosome-sized footprint midpoints occupied virtually all possible positions along the sequences, overriding observed sequence-dependencies on native fibers. Next, we performed visual inspection of individual sampled fibers before and after remodeling (**Figure 2C-D**). Across both fiber types, we observed expected hallmarks of SNF2h remodeling, including the formation of what appear to be evenly-spaced nucleosomal arrays. Importantly, several aspects of our remodeling data recapitulate existing knowledge of how SNF2h binds and remodels mononucleosomes: for instance, remodeling did not substantially impact the estimated numbers of nucleosomes per template, consistent with ISWI remodelers predominantly sliding, not evicting or loading nucleosomes (**Supplementary Table 2**), and the precatalytic condition (SNF2h(-)ATP) yielded larger footprints on average but little change in preferred nucleosome positions on templates, consistent with the HSS domain of SNF2h interrogating DNA flanking the nucleosome (Grüne et al., 2003; Hota et al., 2013; Leonard and Narlikar, 2015).

**Figure 2:**
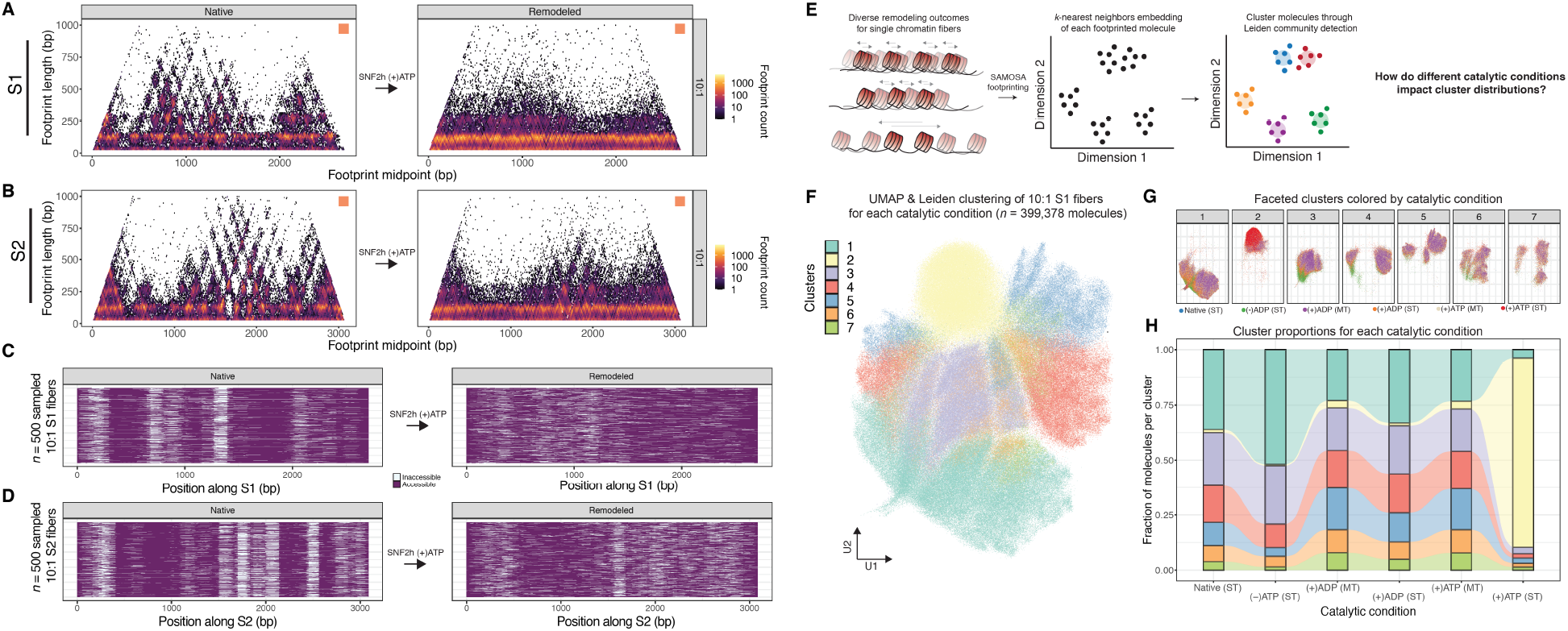
SAMOSA-ChAAT reveals chromatin remodeling outcomes at single-fiber resolution. **A-B.)** Footprint length versus footprint midpoint horizon plots capture bulk outcomes of SNF2h remodeling of 10:1 targeted density S1 fibers (**A**) and S2 fibers (**B**). In both cases, remodeling relaxes bulk sequence preferences for nucleosome assembly. **C-D.)** Sampled single-molecule data for 10:1 S1 (**C**) and S2 (**D**) fibers before and after single-turnover SNF2h remodeling. Remodeling translocates nucleosomes from preferred starting positions, leading to the formation of nucleosome arrays. **E.)** Schematic representation of a computational approach to infer how remodeling alters fiber structure, by using a combination of *k*-nearest neighbors visualization (UMAP) and clustering (Leiden) to define the ‘state-space’ of possible chromatin accessibility patterns. **F.)** UMAP visualization and Leiden clustering (colors) of all 10:1 fibers from six different catalytic condition experiments. **G.)** UMAP visualization faceted by cluster type, colored by catalytic condition. **H.)** Alluvial plots visualize shifts in cluster distribution across each catalytic condition.

We next hypothesized that single-molecule data might offer insight into SNF2h-catalyzed ‘transitions’ between thermodynamically-favorable versus translocated nucleosome positions. To address this, we formalized the UMAP and Leiden clustering methods into a pipeline for visualizing and clustering SAMOSA remodeling data across multiple catalytic conditions (**Figure 2E**), and applied this to native and remodeled 10:1 S1 fibers. UMAP visualization and Leiden clustering (**Figure 2F**) of 10:1 S1 fibers across all catalytic conditions resulted in seven individual clusters. These seven clusters were nonuniformly distributed across each catalytic condition, suggesting that our unsupervised clustering approach can distinguish fiber types on the basis of remodeling condition (**Figure 2G**). Visualizing these distributions as alluvial plots (**Figure 2H**) demonstrates the potential of our approach; by quantifying the relative changes in cluster usage over different conditions (*e.g*. increase in cluster 2 representation under saturating SNF2h remodeling conditions), our data enable a comprehensive view of heterogeneous state spaces. These data and analyses demonstrate the ability of our platform to study and potentially model dynamic changes in chromatin fiber structure at high-throughput and at single-molecule resolution.

### SNF2h does not preferentially clamp trinucleosomes on individual fibers

Our data provide a unique opportunity to distinguish between the ‘clamping’ and ‘length-sensing’ models of SNF2h remodeling (**Figure 3A**). To begin testing these two models, we queried how remodeled fibers were structured at the level of trinucleosomes. Specifically, we computed the following: for every series of three successive footprints on each molecule, we determined the distance between the second and third footprint midpoints (‘*d*_2_’) and plotted this as a function of the distance between the first and second footprint midpoints (‘*d*_1_’; **Figure 3B**). The simplest prediction of the clamping model is that the distances between three successive remodeled nucleosomes should cluster around a fixed length, with unequal distances resulting from trinucleosomes where nucleosome n3 is positioned distantly enough to evade clamping (**Figure 3C**). Conversely, if SNF2h moves nucleosomes via length-sensing, we expect to observe weak correlation between *d_1_* and *d_2_*, with remodeling i.) increasing the correlation between these distances as a function of increasing nucleosome density, and ii.) decreasing the average distance between nucleosomes as a function of density (**Figure 3D**).

**Figure 3:**
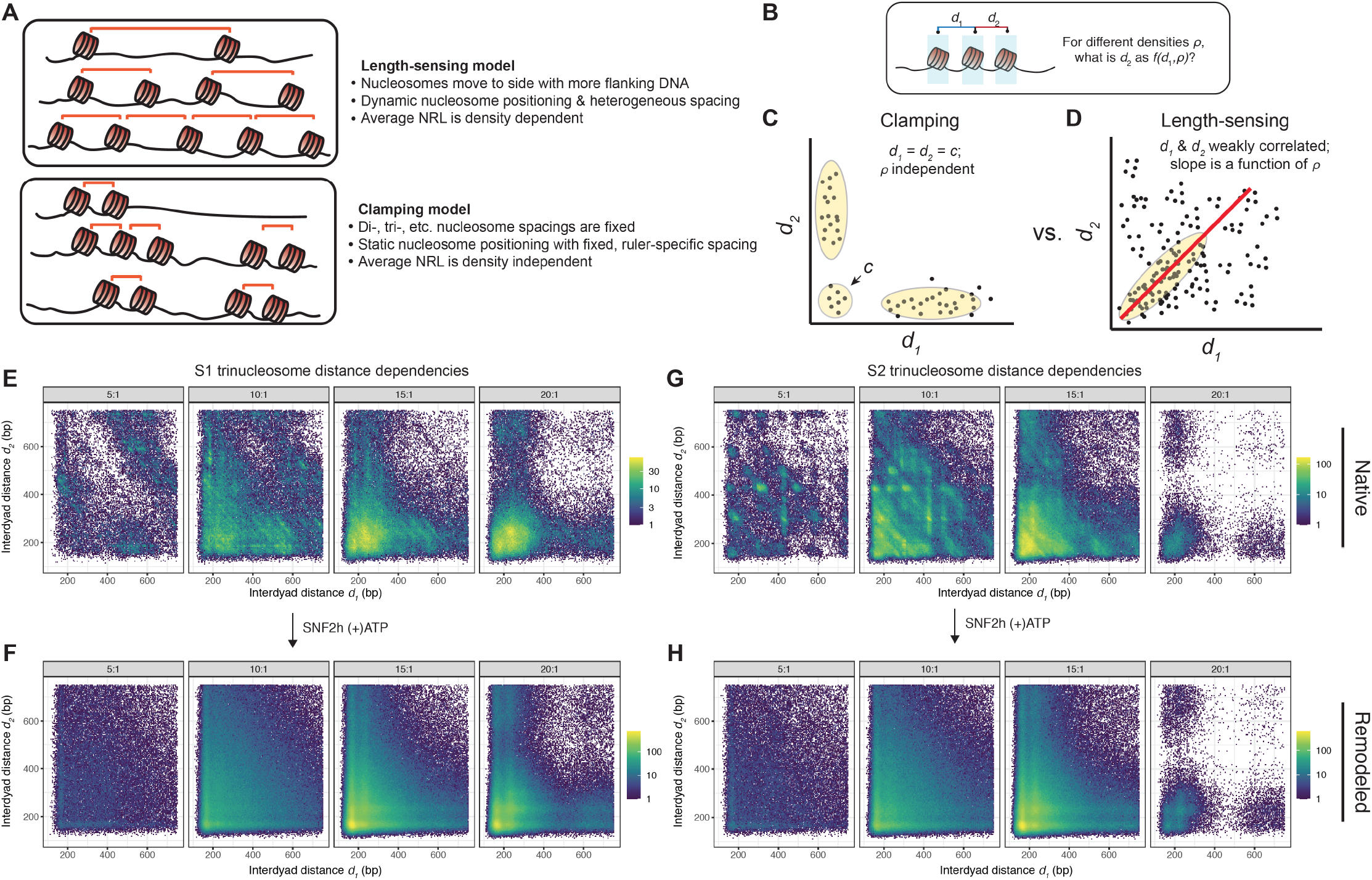
SNF2h does not fix trinucleosomal spacings on individual chromatin fibers. **A.)** Schematic of two competing models for how SNF2h creates regular nucleosome arrays; the ‘length-sensing’ model suggests that array regularity is a steady-state product of flanking DNA sampling by the remodeler and subsequent translocation of nucleosomes towards sides with longer flanking DNA length. The ‘clamping’ model suggests that SNF2h clamps nucleosomes to fixed distances. **B.)** We constructed an analysis to determine how SNF2h remodels trinucleosomes on individual fibers by testing for correlation between the distances between successive nucleosomes. **C.)** Schematic of expected results of the trinucleosome distance analysis if SNF2h acts according ot the ‘clamping’ model. **D.)** Schematic of expected results of the trinucleosome distance analysis if SNF2h acts according to the ‘length-sensing’ model. **E-F.)** Scatter plots of the resulting correlations, for unremodeled S1 (**E**) and remodeled S1 (**F**) fibers. Remodeling diversifies possible internucleosomal distances, as evidenced by the loss of ‘patchy’ sequence-dependent paired nucleosome distances upon remodeling. **G-H.)** As in **E-F.)**, but for unremodeled S2 fibers (**G**) and remodeled S2 fibers.

Trinucleosomal distance scatterplots for unremodeled and remodeled S1 and S2 fibers are visualized in **Figure 3E-H**. In all cases, SNF2h remodeling had a clear visual effect on pairwise distance distributions, converting sequence-programmed distance-dependencies (shown as ‘patches’ of enrichment in unremodeled fibers) into a random configuration of successive distances. The degree to which remodeling impacted the observed correlation between distances varied across different density preparations and different sequences (**Supplementary Figure 5A-B**). In some cases, remodeling altered the sign of the correlation, though this was not necessarily dependent on starting chromatin density. On first glance, our results seem evocative of the clamping model, as fixed distances appear more frequently at higher densities. Comparisons of remodeled scatter plots (**Figure 3F,H**) with unremodeled scatter plots (**Figure 3E,G**), however, reveal that much of this signal is in fact encoded prior to remodeling, and is reliant on high nucleosome density. In reality, SNF2h remodeling products do not display density-independent trinucleosomal spacings, nor do they display favored dinucleosomal spacings (**Supplementary Figure 5C-D**). Instead, our data are consistent with SNF2h translocating nucleosomes towards the longer linker DNA without clamping di- and tri-nucleosomes on individual fibers.

### Density-dependent SNF2h remodeling generates chromatin fibers with a range of regular spacings

To further discriminate between existing models of SNF2h activity, we carried out a second single-molecule analysis—directly detecting array regularity on individual fibers through autocorrelation (**Figure 4A**). We previously demonstrated that single-molecule autocorrelograms and unbiased clustering could be used to simultaneously ascertain regularity and NRL directly from single-molecule data (Abdulhay et al., 2020). We applied this analysis to all footprinted native and remodeled fibers, and then clustered molecules on the basis of similar autocorrelograms, arriving at 8 distinct S1 fiber clusters (average signals shown in **Figure 4B**). These clusters classified footprinted molecules by increasing average distance between nucleosomes across entire single DNA templates, simultaneously capturing molecules with consistent NRLs (*e.g*. clusters 1 - 3), and molecules where a regular pattern was not detected (clusters 7-8; likely owing to hyper- or hypomethylation of the fiber). To ascertain how cluster usage differed as a function of nucleosome density, we visualized cluster enrichment as a heatmap of effect sizes stratified by density (**Figure 4C**), as well as stacked bar graphs capturing the absolute abundance of each cluster as functions of density and remodeling (**Figure 4D**).

**Figure 4:**
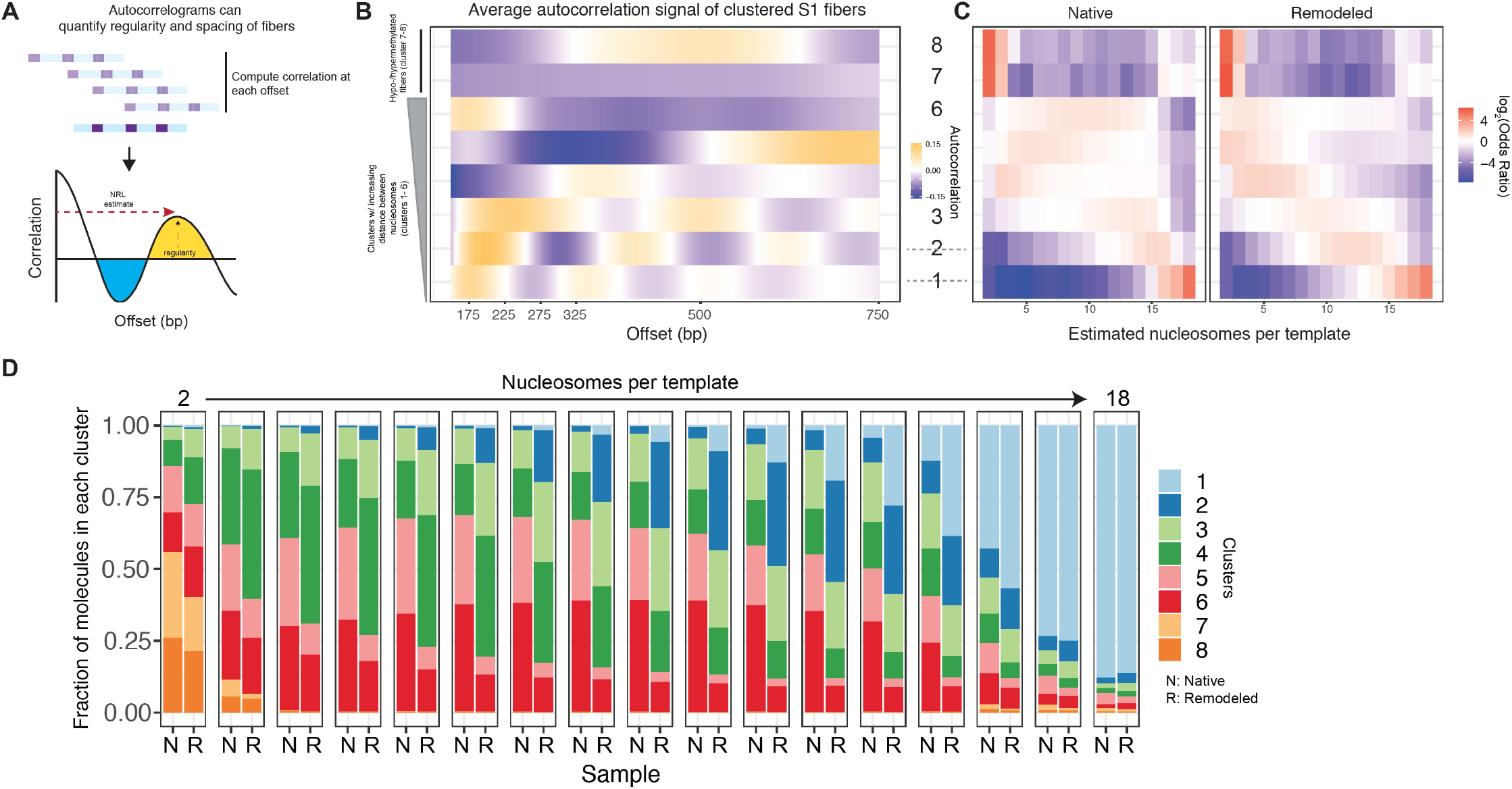
Autocorrelation analyses reveal that chromatin density influences the heterogeneous reaction outcomes of SNF2h remodeling. **A.)** Single-molecule autocorrelation can be used to estimate relative spacing and regularity of single, footprinted chromatin fibers. A schematic of how the autocorrelation function is calculated is illustrated here. **B.)** Results of Leiden clustering of all native and remodeled fiber autocorrelograms. Leiden clustering results in 8 clusters, the first six of which have been classified on the basis of increasing average distance between nucleosomes, and the latter two of which appear to harbor highly hypo- or hyper-methylated fibers. **C.)** Fisher’s exact test enrichment results (log-transformed odds ratios) surveying the relative enrichment and depletion of clusters at each chromatin density (# of nucleosomes per template). Separation of results between ‘Native’ and ‘Remodeled’ classes demonstrates the contribution of statistical nucleosome positioning in unremodeled samples, while illustrating how density can affect fiber state distributions upon remodeling. **D.)** A stacked bar-chart representation of the relative abundance of each fiber-type, stratified by estimated chromatin fiber density; N: native, R: remodeled.

Importantly, our method allowed us to visualize results for both native and remodeled fibers to account for the random formation of nucleosome arrays by statistical positioning downstream of free DNA template ends (Kornberg and Stryer, 1988). Most prior biochemical reactions have been studied at higher nucleosome densities (equivalent of 18 nucleosomes per S1 template) used in our studies. The products of SNF2h remodeling at these densities appear less heterogeneous than the products at lower densities (**Figure 4C,D**). However, at these densities the starting architecture of fibers is also less heterogeneous than at lower densities due to the effects of statistical positioning. More broadly, these results illustrate how nucleosome density influences the state distribution of remodeling outcomes. Even at relatively low fiber densities (*e.g*. 5 – 7 nucleosomes per template), SNF2h remodeling generates a distribution of regular fibers of various predicted NRLs.

To better quantify this distribution before and after remodeling, we computed the probability of observing fibers in each cluster at each nucleosome density, and calculated a log-odds ratio comparing probability distributions for remodeled versus native fibers (**Supplementary Figure 6**). These plots capture the complexity of the ‘state space’ of SNF2h remodeling: remodeling increases the representation (shown in red) of multiple fiber types at multiple densities, with underlying nucleosome density shaping the steady-state distribution of fiber structures. These results, which extend to a completely different primary DNA sequence (**Supplementary Figure 7**), provide further evidence for the length-sensing model of SNF2h activity, and suggest two fundamental properties of remodeling by the SNF2h enzyme: array formation and spacing is influenced by underlying nucleosome density, and reaction outcomes are highly heterogeneous at all densities when the starting nucleosome architectures are explicitly considered. Importantly, these *in vitro* results are inconsistent with the clamping model.

### SNF2h remodeling modulates motif site exposure frequency in a nucleosome density and sequence-dependent manner

How ATP-dependent chromatin remodelers and TFs collaborate in *trans* to access the chromatinized genome is an area of ongoing investigation. We reasoned that our dataset would allow us to ask how nucleosome density, primary sequence, and nucleosome sliding activities synergize to regulate the accessibility of specific sequence motifs. Sequences S1 and S2 collectively harbor three predicted murine 17 bp Ctcf binding motifs (referred to here as S1.1, S2.1, and S2.2), and the accessibility of these motifs *in vivo* depends to various extents on SNF2h activity (Barisic et al., 2019). We aimed to define how SNF2h activity may increase or decrease the nucleosome occupancy likelihood of these motifs (**Figure 5A**). To do so, we examined the relative methyltransferase accessibility of every possible 17-mer along our sequences before and after remodeling and computed a log-odds ratio that captures relative increases (in red) and decreases (in blue) in motif accessibility as a function of sequence position, chromatin density and SNF2h remodeling (**Figure 5B**; S1 fold changes from - 10.5-fold decrease to 63.6-fold increase; S2 fold changes from 25.8-fold decrease to 60.1-fold increase). We highlight two features of SNF2h activity resulting from this analysis: first, remodeler-dependent motif exposure *in vitro* is influenced by both starting nucleosome positions and nucleosome density; second, the ability of SNF2h to create accessibility at particular Ctcf sites (**Figure 5C-E**; specific fold changes noted inline) varies substantially, as remodeling differentially impacts the relative accessibility of sites in a sequence- and density-dependent manner (*e.g*. 3.76-fold increase after remodeling of 8:1 density templates for S1.1; 1.07-fold decrease after remodeling 3:1 density templates for S2.2). The maps presented here provide an essential starting point for modeling how the coupled influences of sequence, nucleosome density, and nucleosome sliding activities quantitatively increase or decrease the probability of site exposure on a population of templates.

**Figure 5:**
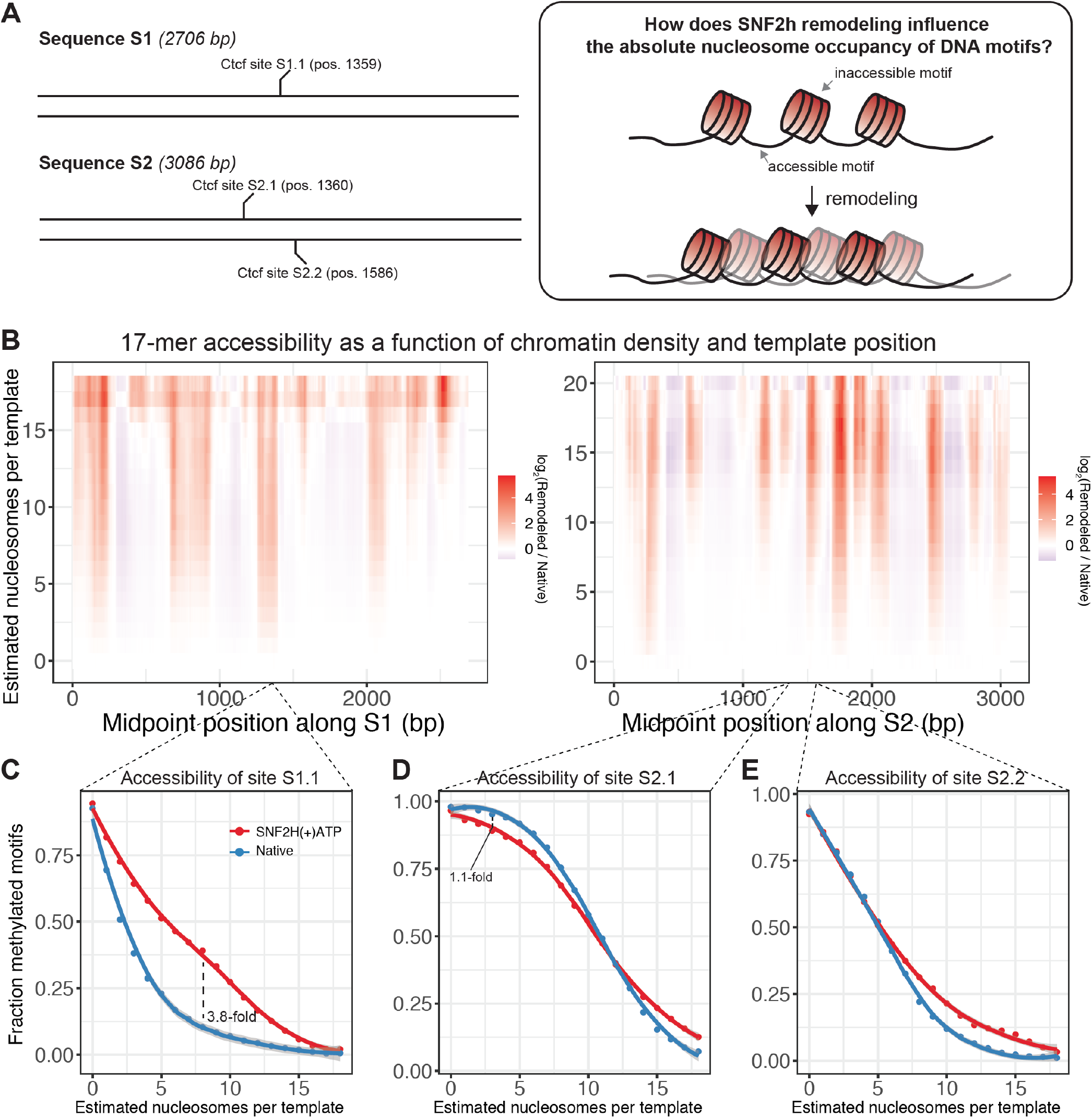
SNF2h remodeling influences motif site exposure in a nucleosome-density and sequence-dependent manner. **A.)** Sequences S1 and S2 collectively contain 3 matches to the canonical 17 bp CTCF / Ctcf binding site (referred to here as S1.1, S2.1, and S2.2). We sought to determine how SNF2h remodeling could influence the probability that these 17 bp sites might be accessible using methylation as a proxy. **B.)** Density vs. sequence heatmaps relating the log_2_ fold change in probability of accessibility of every possible 17 bp motif along sequences S1 and S1. Regions in red represent the 17 bp sites where remodeling increases the probability of methylation; conversely, regions in blue are sites where remodeling decreases the probability of methylation. Accessibility was calculated as a binary variable gated on whether > 90% of the 17 bp motif was methylated on an individual molecule. **C.-E.)** Fractional accessibility as a function of nucleosome density for CTCF / Ctcf site S1.1 (**C**), site S2.1 (**D**), and site 2.2 (**E**) for native fibers (blue) and remodeled fibers (red). Specific values for fold change increases (**C**) or decreases (**D**) highlighted inline.

### SNF2h loss *in vivo* leads to bidirectional, domain-specific shifts in chromatin fiber structure

Substantial prior work has demonstrated that SNF2h-containing complexes can open and repress chromatin accessibility (Barisic et al., 2019; Fyodorov et al., 2004; Wiechens et al., 2016; Xiao et al., 2001), but how these regulatory modes are deployed *in vivo* remains unclear. We speculated that length-sensing by SNF2h may enable these regulatory modes, and performed *in vivo* experiments to test this hypothesis. In mammalian cells, SNF2h (encoded by *SMARCA5 / Smarca5*) is one of two possible catalytic subunits for several ISWI remodeling complexes, each with specific genome-wide localization and activity (Badenhorst et al., 2002; Ito et al., 1997; Varga-Weisz et al., 1997). Importantly, SNF2h is nonessential in murine embryonic stem cells (mESCs), offering a unique opportunity to study how steady-state chromatin fiber structure *in vivo* is impacted by the addition of SNF2h in *trans* (Barisic et al., 2019). We applied an improved, *in situ* version of the SAMOSA protocol (**Supplementary Figure 8**) to footprint feeder-cultured mESCs devoid of SNF2h (*Smarca5^-/-^* mESCs; *i.e*. ‘knockout’), knockout cells expressing a wildtype copy of the SNF2h protein (*i.e*. ‘addback’) (Barisic et al., 2019), and wildtype feeder-free E14 mESCs (*i.e*. ‘E14’). Across all cell lines and including biological replicates, we sequenced 1.66E7 individual fibers, the equivalent of ~9X haploid coverage of the mouse genome. We used these data to ask (**Figure 6A**): i.) how does SNF2h-loss impact the distribution of fiber structures genome-wide; and ii.) where do SNF2-mediated structural changes occur across the mESC epigenome?

**Figure 6:**
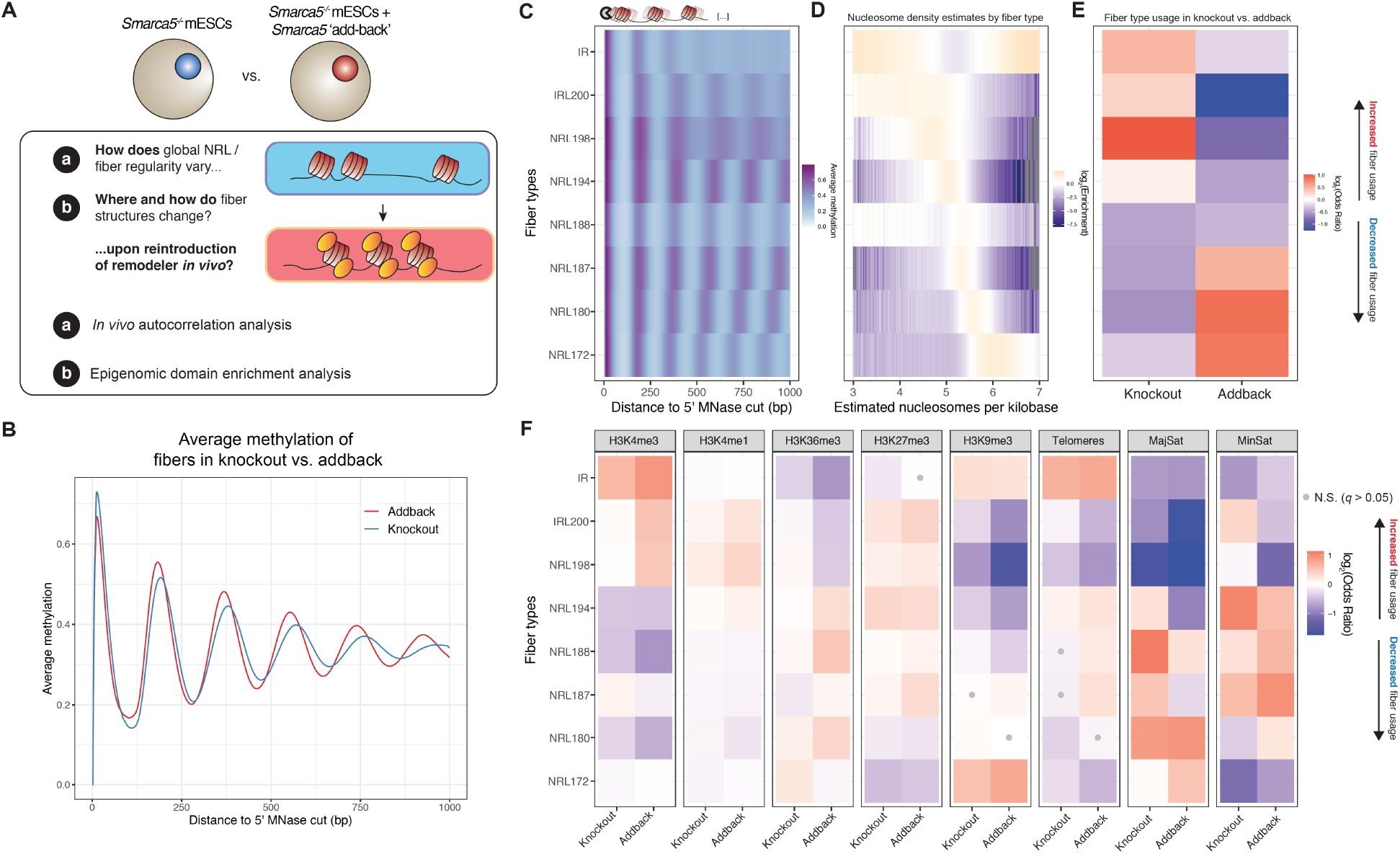
Mapping the *in vivo* consequences of SNF2h remodeling in murine embryonic stem cells at single fiber resolution. **A.)** Overview of *in vivo* experimental design. We performed *in vivo* SAMOSA footprinting in mESCs devoid of SNF2h (*Smarca5^-/-^*; ‘knockout’ cells) and mutant mESCs overexpressing *Smarca5* cDNA (addback cells). We then used the resulting single-molecule data to determine how the addition of SNF2h impacts chromatin fiber architecture globally and at specific epigenomic domains. **B.)** Average single-molecule methylation patterns in knockout cells (blue) and addback cells (red). Reintroduction of SNF2h decreases the overall nucleosome repeat length (NRL) on individual fibers. **C.)** Leiden clustered single-molecule autocorrelograms stratify the mESC epigenome into eight distinct clusters (*i.e*. ‘fiber types’), two ‘irregular’ clusters and six regular clusters arranged here in decreasing NRL. **D.)** Single-molecule nucleosome density estimates for molecules in each of the six clusters, colored by increasing mean nucleosome density. Calculations were made with respect to the background frequency of fiber density estimates. **E.)** Global enrichment of fiber usage plotted as a log-transformed Fisher’s Exact odds ratio of fiber types in knockout versus addback cell lines. Knockout cells are enriched for irregular and long NRL fibers, and the addition of SNF2h leads to the depletion of long NRL / irregular fiber types and enrichment of short NRL fibers in addback cells. **F.)** Differences in fiber type composition across varying epigenomic domains in knockout versus addback cells. Actively transcribed regions are enriched for lower-density fibers, and heterochromatic regions are enriched for higher-density fibers. Enrichments are calculated as log-transformed Fisher’s Exact odds ratios. Tests that are not significant (*q*-value threshold > 0.05) are marked with a grey dot.

We first inspected the average methylation profile of footprinted fibers from knockout and addback cells (**Figure 6B**); consistent with previous results, we found that knockout cells had globally longer NRLs compared to addback cells (Barisic et al., 2019). We then clustered single-molecule autocorrelograms for all data falling within one of eleven different genomic domains. As previously observed in K562 cells (Abdulhay et al., 2020), this analysis resulted in clusters that stratify the genome on the basis of array regularity and NRL (**Figure 6C**). Our unsupervised approach yielded 8 clusters—six regular clusters ranging in NRL from ~172 bp to ~198 bp, and two irregular clusters with weak nucleosome phasing (IRL200) or undetected phasing (IR). These clusters were tightly associated with the underlying nucleosome density of clustered fibers (**Figure 6D**). Knockout cells were globally enriched for irregular IRL200 / IR clusters and low-density NRL198 fibers, while addback cells were conversely enriched for higher density, short NRL fiber types (**Figure 6E**), consistent with the pattern observed on bulk averages. These patterns were highly reproducible, both visually (**Supplementary Figure 9A**) and quantitatively (**Supplementary Figure 9B**).

In previous work, we observed that epigenomic domains were highly heterogeneous with respect to chromatin fiber usage (Abdulhay et al., 2020). In light of this heterogeneity, we sought to determine whether SNF2h loss would consistently impact fiber usage patterns across the epigenome, in line with ‘clamping’ activity *in vivo*. To test this, we examined fibers falling into one of nine different epigenomic domains (H3K4me3, H3K4me1, H3K36me3, H3K27me3, H3K9me3, telomeric sequence, major satellite, and minor satellite), and computed the relative enrichment and depletion of fiber types within each domain in knockout and addback cells (**Figure 6F**; reproducibility shown in **Supplementary Figure 9C-D**). Intriguingly, we found that the reintroduction of SNF2h in addback cells had domain-specific effects: at predicted active promoters, for instance, the addition of SNF2h leads to increased representation of ‘irregular’ and long NRL fibers; at predicted H3K36me3 regions, SNF2h increased the representation of intermediate-length NRL fibers; finally, at typically unmappable heterochromatic major and minor satellite sequences, the addition of SNF2h led to increased representation of short NRL fiber types, consistent with SNF2h condensing chromatin fibers in this context. These results demonstrate that SNF2h remodeling outcomes are domain-specific, suggest that domain-level nucleosome density impacts SNF2h remodeling, and provide further evidence against ISWI clamping *in vivo*.

### SNF2h tunes heterogeneous fiber usage patterns to specify bulk chromatin accessibility

To further explore how SNF2h might differentially regulate specific sites *in vivo*, we re-analyzed paired ATAC-seq datasets from knockout and wildtype mESCs (**Supplementary Figure 10A**) (Barisic et al., 2019). This allowed us to define genomic sites that (directly or indirectly) depend on SNF2h to either open or close chromatin in cell populations. We then extended our enrichment analyses to examine how single-fiber usage patterns are modulated by SNF2h for these sets of loci. Comparing knockout and addback fibers across these sites, we find that sites dependent on SNF2h to remain closed demonstrate a subtle increase in the relative abundance of regular, long NRL arrays, while sites that depend on SNF2h to create accessibility have an increased representation of irregular fibers (**Supplementary Figure 10B**). These fiber usage distributions are highly heterogeneous (**Supplementary Figure 10C**), hinting that SNF2h tunes, rather than specifies, fiber usage patterns to change bulk chromatin accessibility.

We next focused specifically on the transcription factor Ctcf, whose occupancy in mammalian cells depends on SNF2h activity (Barisic et al., 2019; Wiechens et al., 2016). Specifically we examined, across both cell lines, footprinted chromatin fibers containing known gold-standard Ctcf bound sites in the mESC epigenome (Yue et al., 2014), as well as fibers harboring motif matches drawn randomly from the approximately 2.9E6 Ctcf sites found across the murine genome (*i.e*. unoccupied Ctcf sites) (Vierstra et al., 2020). Averaging signal over all of these fibers centered at predicted Ctcf motifs, we recapitulated previous findings, observing that Ctcf site accessibility is dramatically impacted by the loss of SNF2h *in vivo*, although SNF2h-loss does not lead to complete occlusion of factor binding sites (**Figure 7A**; compare with unbound Ctcf sites). These results were orthogonally corroborated by ATAC fragment analyses at these same sites.

**Figure 7:**
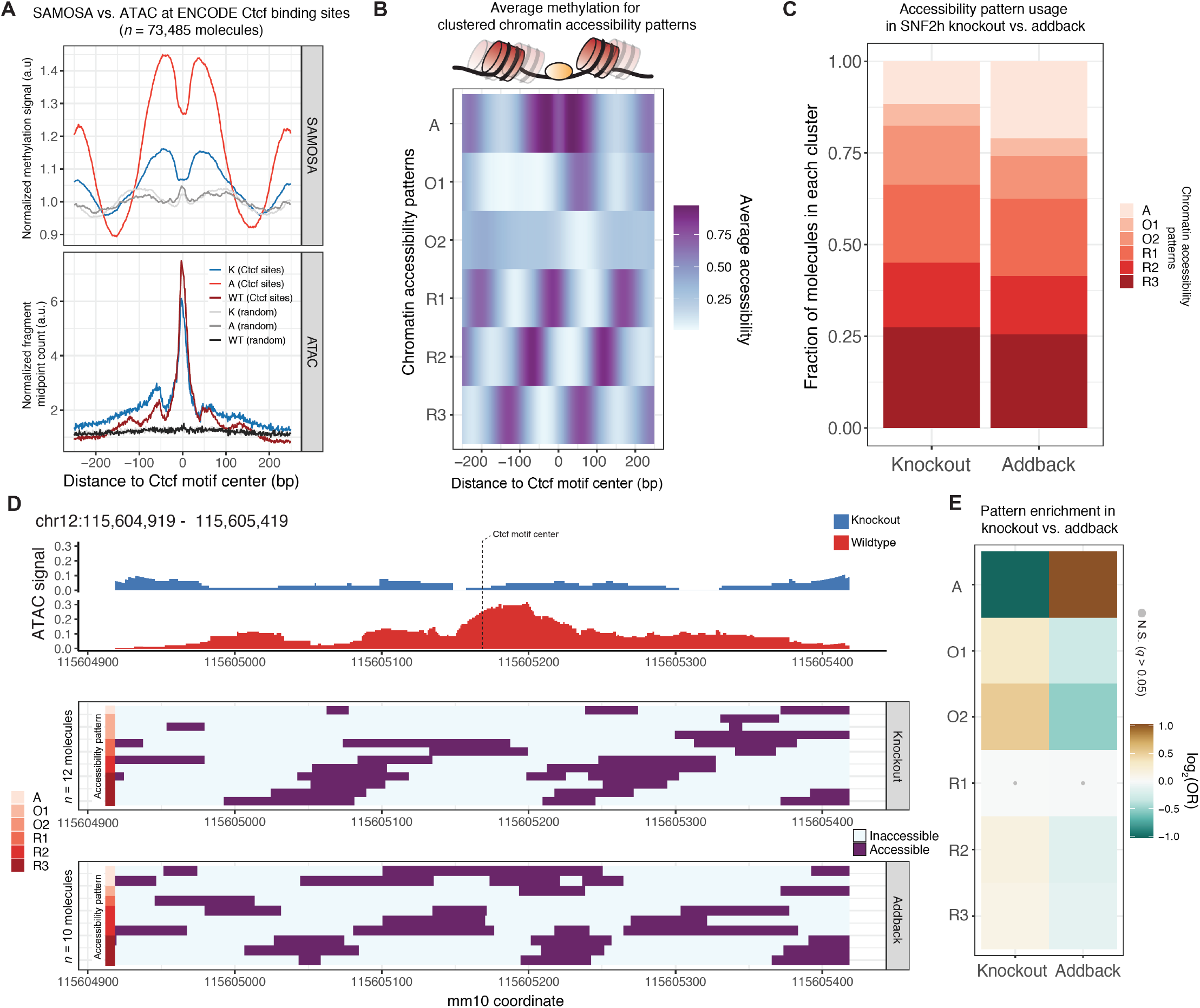
Exploring SNF2h-mediated chromatin closing and opening by integrating ATAC-seq and SAMOSA data. **C.)** Line plots for normalized SAMOSA (top) and ATAC-seq (bottom) signal at ENCODE-defined Ctcf sites and unbound Ctcf motif matches in the mESC epigenome. **D.)** Average accessibility patterns for clustered Ctcf site occupancy patterns. Clusters can be broken down into three regular nucleosome occupancy patterns (R1 – R3), two irregular occupancy patterns (O1-O2), and one accessible cluster (A). **E.)** Stacked bar chart representation of occupancy pattern cluster distribution in knockout vs. addback cells. **F.)** Single-site representation of a Ctcf site where bulk chromatin accessibility is lost in the knockout genome. Single-molecules covering this site are shown below and are labeled with respective cluster labels. **G.)** Fisher’s exact test results for enrichment (gold) or depletion (green) of occupancy patterns in knockout vs. addback cells. Tests that are not significant following *q*-value correction (*q* < 0.05) are marked with a grey dot.

We next quantified the impact of SNF2h remodeling on Ctcf occupancy patterns on single-molecules. To do this, we aggregated all single-molecule observations within a 500 nucleotide window surrounding bound sites and clustered all molecules to determine nucleosome occupancy patterns immediately surrounding Ctcf binding sites. Our analyses yielded 6 primary patterns of nucleosome occupancy over these motifs (**Figure 7B; Supplementary Figure 11**), including two irregular, or ‘offset’ nucleosome occupancy patterns (clusters O1 and O2), three nucleosome occluded patterns harboring well-phased nucleosomes in various registers (clusters R1 – R3), and a TF accessible cluster of molecules, a subset of which display unmethylated footprints potentially capturing direct Ctcf-DNA interactions (cluster A) (**Supplementary Figure 11**). Cluster usage was highly heterogeneous across knockout and addback cells (**Figure 7C**), a finding that was underscored by analysis of a single SNF2h-dependent Ctcf site (12X coverage in knockout; 10X coverage in addback) in the murine genome (**Figure 7D**). At this site, fibers of each type were observed, demonstrating extensive heterogeneity in fiber patterns at a single, SNF2h-regulated locus. In line with this, quantification of differential cluster usage across knockout and addback cells revealed a modest depletion (1.43-fold for cluster O2) for offset clusters and modest enrichment (2.04-fold for cluster A) for the accessible cluster in addback cells. These distributions suggest that SNF2h again tunes chromatin accessibility at these sites through continuous nucleosome sliding, and not through programmed clamping of nucleosomes; we speculate that this sliding is in turn regulated by local nucleosome density, which controls the relative amount of extranucleosomal flanking DNA available to allow SNF2h catalysis (Leonard and Narlikar, 2015).

## DISCUSSION

### Dissecting chromatin remodeling outcomes at single-fiber resolution using SAMOSA-ChAAT

Modern chromatin biology sits amid a ‘resolution revolution.’ Advances in cryogenic electron microscopy (‘cryo-EM’) have provided us with near-atomic views of macromolecular chromatin-interacting complexes (Armache et al., 2019; Eustermann et al., 2018; Patel et al., 2019). Complementarily, advances in single-molecule and high-resolution microscopic approaches *in vitro* and *in vivo* have provided new views of dynamic and often heterogeneous chromatin conformations (Blosser et al., 2009; Boettiger et al., 2016; Kim et al., 2021). Finally, advances in high-throughput short-read sequencing have offered near nucleotide-resolution maps of where and how these complexes engage with chromatin genome-wide, across myriad substrates *in vitro*, and even at the resolution of single-cells (de Dieuleveult et al., 2016; Henikoff et al., 2011; Krietenstein et al., 2016; Lai et al., 2018). SAMOSA-ChAAT provides a fourth advance in chromatin resolution—datasets describing the molecularly-resolved activity of chromatin regulators on individual chromatin fibers. Our data and associated computational pipelines offer a new frontier for quantifying dynamic chromatin-associated processes that complements existing high-resolution approaches. In future work, we anticipate using SAMOSA-ChAAT to study post-translationally modified chromatin fibers, as well as fibers undergoing additional dynamic nuclear processes (*e.g*. transcription, replication, loop extrusion).

### SNF2h is a length-sensing, nucleosome-density dependent chromatin remodeler

Chromatin remodelers regulate nucleosome spacing *in vitro* and *in vivo*, but the question of how chromatin remodelers like SNF2h space nucleosomes on individual fibers remains open. Using SAMOSA-ChAAT, we performed the first (to our knowledge) single-fiber-resolution footprinting experiments on reconstituted, remodeled, murine genomic templates of varying nucleosome density. Our *in vitro* results highlight three key properties of SNF2h remodeling: first, remodeling outcomes are highly heterogeneous and largely ablate sequence-programmed nucleosome positions, consistent with prior findings that SNF2h remodeling rates are insensitive to nucleosome stability, and implying that remodelers can override intrinsic DNA driven nucleosome positioning (Partensky and Narlikar, 2009; Zhang et al., 2011); second, remodeling products bear little evidence of so-called ‘clamping,’ as the final distributions of internucleosomal distances and single-fiber nucleosome arrangements catalyzed by SNF2h remodeling vary as a function of underlying chromatin density; third, both primary sequence and nucleosome density contribute to whether SNF2h increases or decreases DNA site accessibility, which we demonstrate using the cognate binding motif for the CTCF transcription factor (**Figure 8A**). Given our results, we propose an updated model for SNF2h action on chromatin fibers. *In vitro*, SNF2h does not program fixed IDs; instead, length equalization is a steady-state outcome of moving nucleosomes in the direction of the longer linker DNA, consistent with a length-sensing model for SNF2h activity.

**Figure 8:**
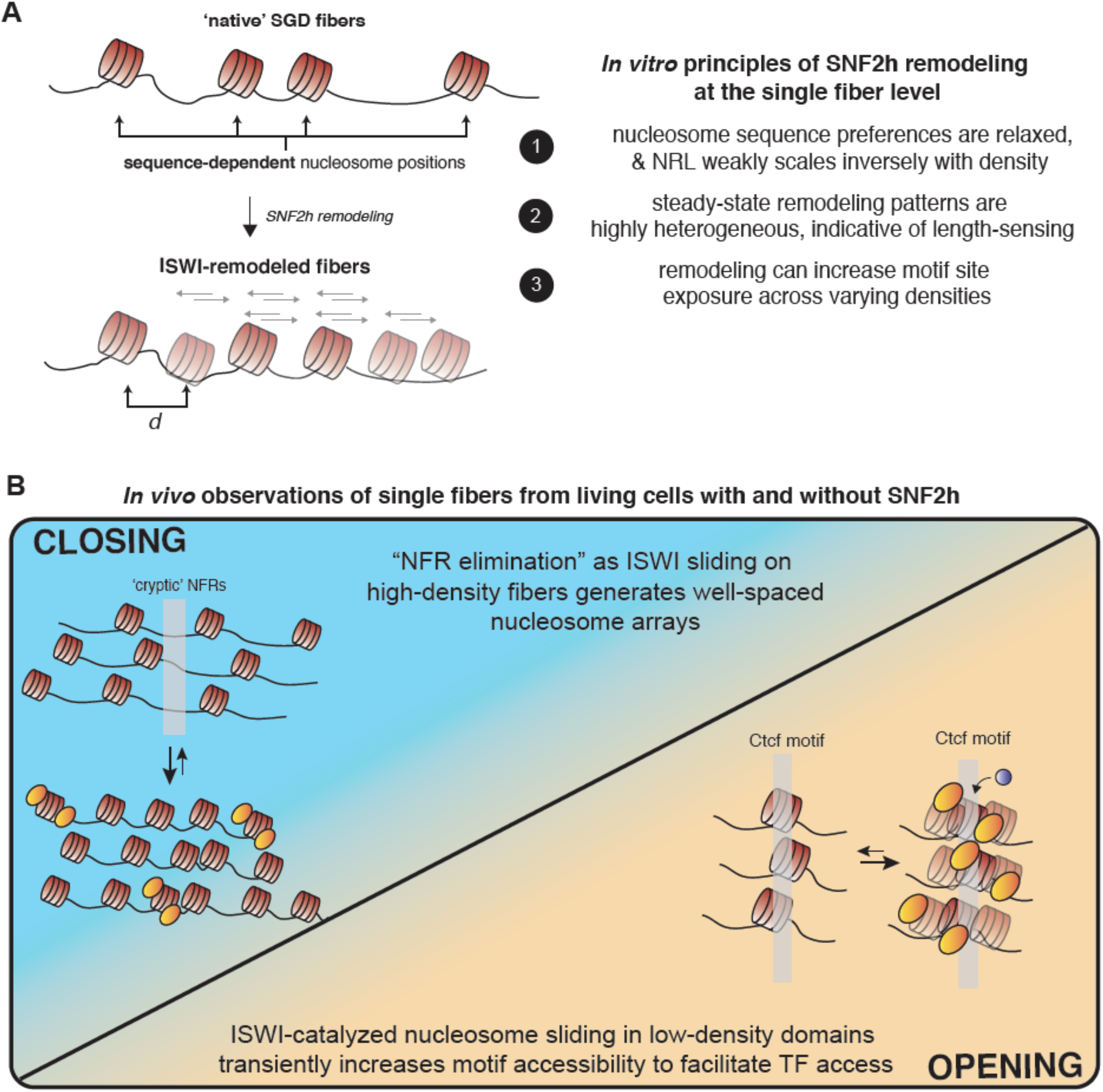
A model of SNF2h-mediated chromatin regulation based on results of this study. **A.)** *In vitro*, SNF2h acts stochastically, sliding nucleosomes that are pre-positioned by primary DNA sequence to new positions. Reaction outcomes are highly heterogeneous and NRLs created by SNF2h remodeling are weakly inversely proportional to the underlying nucleosome density of fiber substrates—evidence that SNF2h operates through length-sensing of extranucleosomal DNA. Finally, remodeling can increase (or decrease) motif site exposure in a nucleosome-density- and sequence-dependent manner. **B.)** *In vivo*, SNF2h length-sensing can explain two diametrically-opposed regulatory functions. At high-nucleosome-density repressed regions, SNF2h increases the representation of multiple types of regular, short NRL fibers, presumably to facilitate the elimination of cryptic NFRs. At lower-nucleosome-density, accessible *cis*-regulatory elements (*e.g*. CTCF / Ctcf binding sites), SNF2h slides nucleosomes to increase the site exposure frequency of these sites, enabling TF access.

### SNF2h employs distinct regulatory modes to repress and open chromatin *in vivo*

How do the length-sensing properties of SNF2h manifest *in vivo*? All of the SNF2h activities discussed here, including length-dependent sliding (Stockdale et al., 2006; Yang et al., 2006; Zofall et al., 2004), active positioning of nucleosomes downstream of barriers (Krietenstein et al., 2016; Yen et al., 2012), and the formation of well-spaced nucleosome arrays (Fyodorov et al., 2004; Ito et al., 1997) have been noted in previous work, but how these sometimes disparate activities harmonize to impact gene regulation *in vivo* has remained elusive. Our data from living mESCs provide evidence that linker-length-sensing can account for all of these classically-defined activities (**Figure 8B**). At regions where SNF2h is required to maintain heterochromatic structure (*i.e*. regions of relatively high nucleosome density), the remodeler converts irregular and long NRL fibers into well-spaced nucleosome arrays with multiple short NRLs. How well-ordered arrays repress chromatin remains unknown, but it is tempting to speculate that this process either facilitates ‘elimination’ of nucleosome-free regions (NFRs) by preventing cryptic NFR formation (Garcia et al., 2010), by promoting chromatin compaction (Correll et al., 2012), or by generating NRLs particularly suited for phase separation (Gibson et al., 2019). At euchromatic regions where SNF2h is required to create chromatin accessibility (*i.e*. regions with relatively low nucleosome density), sliding is now used by SNF2h to generate ‘disordered’ fibers to increase the site exposure frequency of *cis*-regulatory elements like CTCF binding sites. While our data explain the multiple observations of coupled ISWI activity and TF binding (Badenhorst et al., 2002; Barisic et al., 2019; Wiechens et al., 2016; Xiao et al., 2001), how SNF2h remodeling facilitates other dynamic nuclear processes (*e.g*. transcription, replication, repair, higher-order chromatin folding) remains an exciting question.

### Nucleosome density as an additional substrate cue for targeting chromatin remodeling activity

Our understanding of how sequence-non-specific chromatin remodeling complexes achieve specificity at genomic loci is still developing. Prior work has uncovered myriad remodeler-targeting ‘cues,’ including post-translational histone modifications (Clapier et al., 2001; Dann et al., 2017; Mashtalir et al., 2021), TFs (Barisic et al., 2019; Brahma and Henikoff, 2019; de Dieuleveult et al., 2016), three-dimensional (3D) chromosomal architecture (Barisic et al., 2019; Barutcu et al., 2016; Weber et al., 2021), composition of the nucleosome core particle (Dann et al., 2017; Gamarra et al., 2018; McBride et al., 2020), and, as discussed here, availability of extranucleosomal flanking DNA (Stockdale et al., 2006; Yang et al., 2006; Zofall et al., 2004). Here, we elaborate on the length-sensing model by connecting DNA length-sensing on mononucleosomes to nucleosome density of individual chromatin fibers. At high nucleosome densities, flanking DNA is occluded and ISWI remodeling outcomes are constrained to create populations of evenly-spaced arrays capable of repressing chromatin. These fiber-type distributions are likely further regulated by ISWI complex composition (Eberharter et al., 2001; Hamiche et al., 1999; Ito et al., 1997; Varga-Weisz et al., 1997). At low nucleosome densities, extranucleosomal DNA is abundant and ISWI catalysis is unconstrained. This enables continuous nucleosome sliding, allowing *trans* acting factors to overcome nucleosomal repression of regulatory DNA.

### Study limitations

Our study does have limitations. First, we have restricted our analyses to the ISWI ATPase SNF2h—as discussed above, SNF2h operates in the context of complexes whose additional non-catalytic subunits can modulate SNF2h activity. In future work, we anticipate broadly applying SAMOSA-ChAAT to study how ATP-dependent chromatin remodeling complexes and other chromatin-associated factors alter chromatin fiber structure. Second, our assay inherently captures the accessibility of primary sequence in the context of chromatin, which can be influenced by both primary structure (*e.g*. histone-DNA interactions) and higher-order solution chromatin structures (*e.g*. fiber folding). As such, our accessibility patterns must not be overinterpreted as solely representing nucleosome occupancy, and our assay does not capture the same real-time dynamics as *in vitro* and *in vivo* single-molecule experiments. Finally, like other *in vivo* single-molecule footprinting studies to date (Abdulhay et al., 2020; Lee et al., 2020; Shipony et al., 2020; Sönmezer et al., 2021; Stergachis et al., 2020), our mESC data are necessarily averaged over a multitude of discrete cellular states, including the cell cycle. As chromatin structure changes dramatically as a function of cell cycle progression (Stewart-Morgan et al., 2020), a fraction of the single-molecule patterns we observe *in vivo* must represent known ‘bespoke’ chromatin structures (*e.g.* mitotic chromatin). Future work in synchronized or post-mitotic cells will help discriminate the specificity of observed single-molecule patterns at distinct cell cycle stages.

### Concluding remarks

Many questions remain: how is SNF2h activity constrained? SNF2h-containing complexes require specific flanking DNA lengths to enable catalysis—do other factors act in *trans* to effectively gate flanking DNA lengths and inhibit SNF2h activity? How do SNF2h-containing complexes collaborate with other remodeling families? At actively transcribed gene bodies, for instance, INO80, CHD, and ISWI complexes regulate nucleosome positioning in an evolutionarily-conserved manner, and dissecting the contribution of each remodeler remains important. How might nucleosome density (and, by extension, SNF2h activity) be controlled *in vivo*? In mammals, nucleosome density is likely regulated on a diversity of length scales, ranging from local (*e.g*. ATP-dependent chromatin remodeling; histone chaperones; histone modification; replication, transcription, and repair), to global (*e.g*. genome compartmentalization / phase separation; loop extrusion; subnuclear localization). We envision a regulatory circuit wherein the concentration of core histones can be tuned within large chromatin domains by specific *trans* and *cis*-regulatory elements. This circuit could influence the regulatory outputs of remodeling complexes over long genomic distances, allowing higher-order genome conformation to instruct local interpretation of regulatory DNA.

## METHODS

### Cloning *M. musculus* genomic sites for nucleosome array assembly

Two separate sites within the *M. musculus* reference genome containing CTCF sites were chosen for histone assembly. The CTCF genomic sites will be referred to as Sequence 1 “S1” (chr1:156,887,669-156,890,368, 2712bp) and Sequence 2 “S2” (chr1:156,890,410-156,893,258, 2861 bp). S1 and S2 were PCR amplified (NEBNext^®^ Q5 2X Master Mix) from purified E14 mESC genomic DNA with primers containing homology to a Zeocin-resistance multicutter plasmid backbone as well as dual EcoRV sites for downstream separation of insert from backbone. The plasmid backbone sequence of interest containing homology was prepared with PCR amplification and the remaining parental plasmid was digested away (1 uL DpnI in 1X CutSmart at 37°C for 1 hour). All PCR products were subsequently run out on a 1% Agarose gel and gel purified. After gel purification, standard Gibson Cloning for S1 or S2 inserts plus PCR-amplified/DpnI-digested backbone was performed using NEBuilder^®^ HiFi DNA Assembly Master Mix (New England Biolabs) at 3:1 insert to vector ratio. Transformation was performed with Stellar Competent Cells (Takara) which were thawed on ice. 2 uL of assembly reaction was added to 50 uL competent cells and flicked to mix 4-5 times. The mixture was incubated on ice for 30 minutes, heat shocked at 42°C for 30 seconds, and placed on ice for 2 minutes. 950 uL of SOC media was added to the mixture, and an outgrowth step was performed at 37°C for 1 hour shaking at 1000RPM. The entire mixture was added to pre-warmed Zeocin plates and incubated overnight. Colony PCR was performed to test for insert presence – 8 colonies were selected per site and ran on a 1% Agarose gel. Four colonies containing the insert were selected per sequence and miniprepped overnight in Low Salt LB Broth containing Zeocin (25ug/mL). Plasmids were subsequently Sanger Sequenced (Genewiz) to confirm insert sequence and one clone was selected per site for downstream experiments.

### Preparation of S1 and S2 arrays via Salt Gradient Dialysis

To assemble nucleosomes onto the sequences of interest, the S1 and S2 plasmids were purified using a GigaPrep kit (Qiagen). To isolate the insert, purified plasmids were restriction enzyme digested (S1: EcoRV, ApaLI, XhoI, BsrBI and S2: EcoRV, BsrBI, BssSaI/BssSi-v2, FseI, BstXI, PflFI). Each insert was purified by size exclusion chromatography. Plasmid gigapreps were performed with a dam^+^ *E. coli* strain, - GATC sequences were ignored for downstream analysis of *in vitro* experiments. Initial restriction enzyme tests were performed with the plasmids to confirm proper digestion of the backbone, so that the insert could be purified. Xenopus histones were purified according to previously described methods (Luger et al., 1999), and chromatin was assembled using salt gradient dialysis (Luger et al., 1999), with varying ratios of histone:DNA.

### Enzyme remodeling on *in vitro* oligonucleosome chromatin arrays

S1 or S2 arrays assembled at varying histone:DNA concentrations (50 nM arrays) were remodeled under single-turnover, saturating enzyme conditions (9 uM SNF2h) or under multiple turnover conditions (10 nM SNF2h). All remodeling reactions were performed in 12.5 mM HEPES pH 7.5, 3 mM MgCl_2_, 70 mM KCl, and 0.02% NP-40. Reactions were started with the addition of saturating ATP, ADP (2mM) or no nucleotide and incubated for 15 minutes at room temperature. All reactions were quenched immediately with an equal volume of ADP (34mM) in 1X TE resulting in 25 nM arrays.

### SAMOSA on *in vitro* oligonucleosome chromatin arrays

SAMOSA was performed on remodeled arrays as well as unremodeled arrays and unassembled DNA controls using the non-specific adenine EcoGII methyltransferase (New England Biolabs, high concentration stock 2.5e4 U/mL) as previously described (Abdulhay et al., 2020). For the remodeled arrays, entire reaction volume was methylated with 31.25 U (1.25uL) of EcoGII. For unremodeled arrays, 1000 nM of input was methylated with 2.5uL EcoGII. For the unassembled, naked S1 and S2 DNA, 3ug input DNA was methylated with 5ul of EcoGII. Methylation reactions were performed in a 100uL reaction containing 1X CutSmart Buffer and 1mM S-adenosyl-methionine (SAM, New England Biolabs) and incubated at 37°C for 30 minutes. SAM was replenished to 3.15 mM after 15 minutes. Unmethylated S1 and S2 naked DNA controls were similarly supplemented with Methylation Reaction buffer, minus EcoGII and replenishing SAM, and the following purification conditions. To purify the remodeled and unremodeled DNA, the samples were subsequently incubated with 10uL Proteinase K (20mg/mL) and 10uL 10% SDS at 65°C for a minimum of 2 hours up to overnight. To extract the DNA, equal parts volume of Phenol-Chloroform-Isoamyl was added and mixed vigorously by shaking and then spun (max speed, 2 min). The aqueous portion was carefully removed and 0.1x volume 3M NaOAc, 3uL of GlycoBlue, and 3x volume of 100% EtOH were added, mixed gently by inversion, and incubated either at −80°C for four hours or overnight at −20°C. Samples were spun (max speed, 4°C, 30 min), washed with 500uL of 70% EtOH, air dried, and resuspended in 50uL EB Buffer. Sample concentration was measured by Qubit High Sensitivity DNA Assay.

### Preparation of *in vitro* SAMOSA PacBio SMRT Libraries

The Purified DNA from array and DNA samples was used in entirety as input for PacBio SMRTbell library preparation as previously described (Abdulhay et al., 2020). Briefly, preparation of libraries included DNA damage repair, end repair, SMRTbell ligation, and Exonuclease cleanup according to manufacturer’s instruction. After Exonuclease cleanup and a double 0.45x Ampure PB Cleanup, sample concentration was measured by Qubit High Sensitivity DNA Assay (1uL each). To assess for library quality, samples (1uL each) were run on the Agilent Tapestation D5000 Assay. Libraries were sequenced on Sequel II 8M SMRTcells in-house. *In vitro* experiment data were collected over several pooled 30 h Sequel II movie runs with either a 0.6 h or 2 h pre-extension time and either a 2 h or 4 h immobilization time.

### Cell lines and cell culture

Published SNF2h knockout and re-expression mouse embryonic stem cells were provided under MTA by the Dirk Schübeler Laboratory at FMI (Barisic et al., 2019). Cells were thawed and grown for at least two passages onto CF-1 Irradiated Mouse Embryonic Feeder cells (Gibco A34181). Feeder cells were depleted from mESCs for at least two passages prior to collection for SAMOSA experiments. E14 mESCs were gifted from Elphege Nora Laboratory at UCSF. All cell lines were mycoplasma tested upon arrival, routinely tested, and confirmed negative with PCR (NEBNext^®^ Q5 2X Master Mix). All feeder and mESC cultures were grown on 0.2% gelatin. mESCs were maintained in KnockOut DMEM 1X (Gibco) supplemented with 10% Fetal Bovine Serum (Phoenix Scientific, Lot# BW-067C18), 1% 100X GlutaMAX (Gibco), 1% 100X MEM Non-Essential Amino Acids (Gibco), 0.128mM 2-Mercaptoethanol (BioRad), and 1X Leukemia Inhibitory Factor (purified and gifted by Barbara Panning Lab at UCSF).

### SAMOSA on mESC-derived oligonucleosomes

#### Isolation of nuclei

Nuclei were collected for the *in vivo* SAMOSA protocol as previously described (Abdulhay et al., 2020). Briefly, all nuclei were collected per cell line by centrifugation (300xg, 5 min), washed in ice cold 1X PBS, and resuspended in 1 mL Nuclear Isolation Buffer ((20mM HEPES, 10mM KCl, 1mM MgCl2, 0.1% Triton X-100, 20% Glycerol, and 1X Protease Inhibitor (Roche)) per 5-10e6 cells by gently pipetting 5x with a wide-bore tip to release nuclei. The suspension was incubated on ice for 5 minutes, and nuclei were pelleted (600xg, 4°C, 5 min), washed with Buffer M (15mM Tris-HCl pH 8.0, 15 mM NaCl, 60mM KCl, 0.5mM Spermidine), and spun once again. Nuclei were counted via hemocytometer and either slow frozen or split for each experimental condition (plus or minus EcoGII methylation). To slow freeze nuclei, nuclei were resuspended in Freeze Buffer (20mM HEPES pH 7.5, 150mM 5M NaCl, 0.5mM 1M spermidine (Sigma), 1X Protease Inhibitor (Roche), 10% DMSO) and stored at −80°C.

#### Adenine methylation, MNase digest, and overnight dialysis

To proceed to the modified *in vivo* SAMOSA protocol for direct methylation of nuclei, fresh nuclei were resuspended in Methylation Reaction Buffer (Buffer M containing 1mM SAM). 200uL methylation reactions were performed (10uL EcoGII per 1e6 nuclei) and incubated at 37°C for 30 minutes. SAM was replenished to 6.25mM after 15 minutes. Unmethylated controls were similarly supplemented with Buffer M + SAM, minus EcoGII and replenishing SAM. Samples were spun (600xg, 4°C, 5 min) and resuspended in cold MNase digestion Buffer (Buffer M containing 1mM CaCl2). MNase digestion of nuclei was performed in 200uL reactions and 0.02 Units of MNase was added per 1e6 nuclei (Sigma, micrococcal nuclease from *Staphylococcus aureus*) at 4°C for either 45 minutes or 1 hour. EGTA was added to 2mM to stop the digestion and incubated on ice. For nuclear lysis and liberation of chromatin fibers, MNase-digested nuclei were collected (600xg, 4°C, 5 min) and resuspended in ~250uL of Tep20 Buffer (10mM Tris-HCl pH 7.5, 0.1mM EGTA, 20mM NaCl and 1X Protease Inhibitor (Roche) added immediately before use) supplemented with 300ug/mL of Lysolethicin (Sigma, L-α-Lysophosphatidylcholine from bovine brain) and rotated overnight at 4°C. Dialyzed samples were spun to remove nuclear debris (12,000xg, 4°C, 5 minutes) and soluble chromatin fibers in the supernatant were collected. Sample concentration was measured by Nanodrop and chromatin fibers were analyzed by standard agarose gel electrophoresis.

To generate a naked DNA positive control for downstream analysis, gDNA was extracted from E14 mESCs with Lysis Buffer (10mM Tris-Cl pH 8.0, 100mM NaCl, 25mM EDTA pH 8.0, 0.5% SDS, 0.1mg/mL Protease K) and purified with the following conditions. Methylation reactions were performed as previously stated, with 3ug DNA as input and 5uL EcoGII (125U), followed by a second purification as follows. To purify all DNA samples, reactions were incubated with 10uL of RNaseA at room temperature for 10 minutes, followed by 10uL Proteinase K (20mg/mL) and 10uL 10% SDS at 65°C for a minimum of 2 hours up to overnight. To extract the DNA, equal parts volume of Phenol-Chloroform was added and mixed vigorously by shaking, and spun (max speed, 2 min). The aqueous portion was carefully removed and 0.1x volumes of 3M NaOAc, 3uL of GlycoBlue and 3x volumes of 100% EtOH were added, mixed gently by inversion, and incubated overnight at −20°C. Samples were then spun (max speed, 4°C, 30 min), washed with 500uL 70% EtOH, air dried and resuspended in 50uL EB. Sample concentration was measured by Qubit High Sensitivity DNA Assay.

### Preparation of *in vivo* SAMOSA PacBio SMRT Libraries

Purified DNA from mESCs (methylated, unmethylated, naked DNA positive controls) was used to prepare PacBio SMRT libraries using either the SMRTbell Express Template Prep Kit 1.0 (blunt end ligation) or 2.0 (A/T overhang ligation). For the SNF2h KO and SNF2h WT AB mESC purified SAMOSA samples, a minimum of 500ng up to 1.5ug was utilized as input with SMRTbell Express Template Prep Kit 1.0. For the E14 mESCs, a minimum of ~400ng up to 1.7ug was utilized as input with the SMRTbell Express Template Prep Kit 2.0. The naked DNA E14 positive control was sheared with a Covaris G-Tube (5424 Rotor, 3381xg for 1 minute) and sheared to approximately 10,000 bp. Sample size distribution was checked with the Agilent Bioanalyzer DNA chip. The entire sample was utilized as input for library preparation with the PacBio SMRTbell Express Template Prep Kit 2.0. Briefly, all library preparations included DNA damage repair, end repair, SMRTbell ligation with either blunt or overhang unique adapters, and Exonuclease cleanup according to manufacturer’s instructions. Unique PacBio SMRTbell adapters (100uM stock) were annealed to 20uM in annealing buffer (10mM Tris-HCl pH 7.5 and 100mM NaCl) in a thermocycler (95°C 5 min, RT 30 mins, 4°C hold) and stored at −20°C for long-term storage. After exonuclease cleanup and Ampure PB cleanups (0.45X for 1.0 preparation or 1X for 2.0 preparation), the sample concentrations were measured by Qubit High Sensitivity DNA Assay (1uL each). To assess for size distribution and library quality, samples (1 uL each) were run on an Agilent Bioanalyzer DNA chip. Libraries were sequenced in house on Sequel II 8M SMRTcells. *In vivo* data were collected over several pooled 30 h Sequel II movie runs with either a 0.6 h or 2 h pre-extension time and either a 2 h or 4 h immobilization time.

### SMRT data processing

We applied our method to two use cases in the paper, and they differ in the computational workflow to analyze them. The first is for sequencing samples where every DNA molecule has the same sequence, which is the case for our remodeling experiments on the S1 and S2 sequences, presented in **Figures 1-5**. The second use case is for samples from cells containing varied sequences of DNA molecules, such as the murine *in vivo* follow up experiments presented in **Figures 6-7**. The first will be referred to as homogeneous samples, and the second as genomic samples. The workflow for homogenous samples will be presented first in each section, and the deviations for genomic samples detailed at the end.

#### Sequencing read processing

Sequencing reads were processed using software from Pacific Biosciences. The following describes the workflow for homogenous samples:

1. Demultiplex reads Reads were demultiplexed using lima. The flag ‘ --same’ was passed as libraries were generated with the same barcode on both ends. This produces a BAM file for the subreads of each sample.
2. Generate Circular Consensus Sequences (CCS) CCS were generated for each sample using ccs. Default parameters were used other than setting the number of threads with ‘-j’. This produces a BAM file of CCS.
3. Align subreads to the reference genome pbmm2, the pacbio wrapper for minimap2 (Li, 2018), was run on each subreads BAM file (the output of step 1) to align subreads to the reference sequence, producing a BAM file of aligned subreads.
4. Generate missing indices Our analysis code requires pacbio index files (.pbi) for each BAM file. ‘pbmm2’ does not generate index files, so missing indices were generated using ‘pbindex’.

For genomic samples, replace step 3 with this alternate step 3

3. Align CCS to the reference genome Alignment was done using pbmm2, and run on each CCS file, resulting in BAM files containing the CCS and alignment information.

#### Extracting IPD measurements

The IPD values were accessed from the BAM files and log10 transformed after setting any IPD measurements of 0 frames to 1 frame. Then, for each ZMW, at each base in the CCS (for genomic samples) or amplicon reference (for homogenous samples), for both strands, the log transformed IPD values in all subreads were averaged. These mean log IPD values for the molecule were then exported along with the percentiles of log IPD values across subreads within that molecule.

### Predicting methylation status of individual adenines

#### Predicting methylation in homogenous samples

For homogenous samples dimensionality reduction was used to capture variation in IPD measurements between molecules, and then the reduced representations and IPD measurements were used to predict methylation. For each of S1 and S2, the non-adenine mean log IPD measurements from one unmethylated control sample were used to train a truncated singular value decomposition model. The input measurements had the mean of each base subtracted before training. The Truncated SVD class of scikit-learn was used and trained in 20 iterations to produce 40 components. The trained model was then used to transform all molecules in all samples into their reduced representations. Each resulting component had its mean subtracted and was divided by its standard deviation.

Next, a neural network model was trained to predict the mean log IPD at each base in unmethylated control molecules. The dimension reduced representation of the molecules were provided as input to the model, and the output was a value for each adenine on both strands of the amplicon molecule. The neural network was composed of four dense layers with 600 units each, with relu activation and he uniform initialization. A 50% dropout layer was placed after each of these four layers. A final dense layer produced an output for each adenine in the amplicon reference. The model was trained on a negative control sample using Keras, Adam optimizer, mean square error loss, 100 epochs and a batch size of 128. The trained model was then used to predict the mean log IPD value at all adenines in all molecules in all samples. This prediction was subtracted from the measured mean log IPD to get residuals.

A large positive residual represents slower polymerase kinetics at that adenine than would be expected given the sequence context and molecule and is thus evidence of methylation. To find a cutoff of how large the residual should be to be called as methylated, we assembled a dataset of residuals from an equal proportion of molecules from a fully methylated naked DNA control and an unmethylated control. For each individual adenine a student’s t-distribution mixture model was fit to the residuals using the python package smm (Peel and McLachlan, 2000). A two-component model was fit with a tolerance of 1e-6, and a cutoff was found where that residual value was equally likely to originate from either of the two components. Adenines were then filtered by whether a sufficiently informative cutoff had been found. The three criteria for using the methylation predictions at that adenine in further analysis were: 1) The mean of at least one t-distribution had to be above zero, 2) The difference between the means of the two t-distributions had to be at least X, where X was chosen separately for each amplicon reference but varied from 0.1 to 0.3, and 3) At least 2% of the training data was over the cutoff. These were lenient cutoffs that allowed the methylation predictions at 90+% of adenines to be included in downstream analysis. This was done because the next HMM step accounts for the frequency of methylation predictions in unmethylated and fully methylated control samples, and thus adenine bases where methylation prediction was poor will be less informative of DNA accessibility.

#### Predicting methylation in genomic samples

Methylation prediction was made in a similar fashion for genomic samples, with deviations necessitated by the differences in the data. Unlike in homogenous samples, dimensionality reduction could not be used to capture inter-molecular variation due to varying DNA sequences. Instead IPD percentiles were used as neural network inputs. As described above in Extracting IPD measurements, log IPD percentiles were calculated across all subreads in each molecule separately for each template base. Every 10th percentile from 10th to 90th inclusive, for template bases C, G, and T, were used as neural network input. The other input was the DNA sequence context around the measured base, given for three bases 5’ of the template adenine and ten bases 3’ of the template adenine, one-hot encoded. The neural network was a regression model predicting the measured mean log IPD at that template adenine. The neural network consisted of four dense layers with 200 units each, relu activation, and he uniform initialization. The training data was 5,000,000 adenines each from six different unmethylated control samples. The validation data for early stopping was 5,000,000 adenines from each of two more unmethylated control samples. The model was trained using Keras, Adam optimizer, 20 epochs with early stopping (patience of 2 epochs), and a batch size of 128.

To determine at which adenines the methylation prediction was usefully informative and accurate, we used a second neural network model to predict the IPD residual in a positive control sample from sequence context. Sequence contexts that consistently produced residuals near zero in a positive control would be likely never methylated by EcoGII, or always methylated endogenously. The input to this network was the one-hot encoded sequence context as described above. The output was the measured log IPD with predicted log IPD subtracted. The training data was a fully methylated naked DNA sample of E14. Mean log IPD residuals were calculated using the above trained model. 20,000,000 adenines were used as training data and 10,000,000 as validation data. The neural network consisted of three dense layers of 100 units, relu activation, and he uniform initialization. The model was trained using Adam optimizer for two epochs with a batch size of 128. After examining the output of the trained model on negative and positive controls and chromatin, we settled on a cutoff of 0.6 for the predicted residual in positive control for calling a sequence context as usable for downstream analysis, and a cutoff of 0.42 for the mean log IPD residual for calling an adenine as methylated.

### Predicting molecule-wide DNA accessibility using Hidden Markov Models

#### Predicting DNA accessibility in homogeneous samples

To go beyond individual methylation predictions and predict DNA accessibility along each molecule we applied a Hidden Markov Model (HMM). An HMM model was constructed for each amplicon reference, with two states for every adenine at which methylation was predicted: one state representing that adenine being inaccessible to the methyltransferase, and another representing it being accessible. The emission probabilities were all Bernoulli distributions, with the probability of observing a methylation in an inaccessible state being the fraction of unmethylated control molecules predicted to be methylated at that adenine, and the probability of observing a methylation in an accessible state being the fraction of fully methylated naked DNA control molecules predicted to be methylated at that adenine. 0.5 was added to the numerator and denominator of all fractions to avoid any probabilities of zero. An initial state was created with an equal probability of transitioning into either accessible or inaccessible states. Transition probabilities between adenines were set using the logic that for an expected average duration in a single state of L, by the geometric distribution at each base the probability of switching states at the next base will be 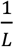. The probability of staying in the same state from one adenine to the next is thus 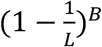, where B is the distance in bases between adenines. The probability of switching to the other state at the next adenine is then 1 minus that value. Different values of the average duration L were tested, and ultimately a value of 1000 bp was used. This is much higher than expected, but has the beneficial result of requiring a higher burden of evidence to motivate switching states and thus minimizes spurious switching.

With the HMM model constructed, the most likely state path was found using the Viterbi algorithm for all molecules in all samples, with the predicted methylation at each adenine provided as the input. Models were constructed and solved using pomegranate (Schreiber, 2018). The solved path was output as an array with accessible adenines as 1, inaccessible as 0, and non-adenine and uncalled bases interpolated.

#### Predicting DNA accessibility in genomic samples

In genomic samples DNA accessibility was predicted in a similar fashion to homogenous, except that the HMM model had to be individually constructed for each molecule due to varying DNA sequences, and rather than empirically measuring the fraction of methylation in positive and control samples at each position, neural networks were trained to predict the fraction of methylation in each from sequence context.

A neural network model was trained to predict the predicted methylation status of adenines in the positive control sample based on sequence context. The output from this model was used to approximate the probability of an adenine in that sequence context getting predicted as methylated if it was accessible to EcoGII. The sample used for training was the same naked DNA E14 methylated sample used to train the positive residual prediction model. Approximately 27,600,000 adenines were used as the training set and 7,000,000 as the validation set. The input was the one-hot encoded sequence context. The neural network consisted of three dense layers of 200 units, relu activation and he uniform initialization. The training output were binary methylation predictions, so the final output of the network had a sigmoid activation and binary cross-entropy was used as the loss. The model was trained with Adam optimizer for seven epochs with the batch size increasing each epoch from 256 to a max of 131,072.

An identical network was trained to predict the predicted methylation status of adenines in the unmethylated negative control samples. The output from this model was used to approximate the probability of an adenine in that sequence context getting predicted as methylated if it was not accessible to EcoGII. This one was trained using adenines combined from four different unmethylated samples, and approximately 28,100,000 adenines were used as the training set and 7,100,000 as the validation set.

The HMM models were constructed in an identical manner to that described above for homogenous samples, except for genomic data an HMM model was constructed for each sequenced molecule individually. States and transition probabilities and observed output were the same. The emission probability of observing methylation at each accessible state was the output of the trained positive control methylation prediction model, and for inaccessible states was the output of the trained negative control methylation prediction model. As with homogenous samples, the HMM was solved using the observed methylation and the Viterbi algorithm.

### Defining inaccessible regions and counting nucleosomes

Inaccessible regions were defined from the HMM output data as continuous stretches with accessibility <= 0.5. To estimate the number of nucleosomes contained within each inaccessible region, a histogram of inaccessible region lengths was generated for each data type (sequence S1, S2, and murine in vivo). Periodic peaks in these histograms were observed that approximated expected sizes for stretches containing one, two, three, etc. nucleosomes. Cutoffs for the different categories were manually defined using the histogram, including a lower cutoff for sub-nucleosomal regions (**Supplementary Figure 12**).

### Processed data analysis

All processed data analyses and associated scripts will be made available at https://github.com/RamaniLab/SAMOSA-ChAAT. Most processed data analyses proceeded from data tables generated using custom python scripts. Resulting data tables were then used to compute all statistics reported in the paper and perform all visualizations (using tidyverse and ggplot2 in R). Below, we describe each analysis in text form, while noting that all code is freely available at the above link.

#### UMAP and Leiden clustering analyses

All UMAP and Leiden clustering analyses were performed using the scanpy package (Wolf et al., 2018). All UMAP visualizations (McInnes et al., 2018) were made using default parameters in scanpy. Leiden clustering (Traag et al., 2019) was performed using resolution = 0.4; clusters were then filtered on the basis of size such that all clusters that collectively summed up to < 5% of the total dataset were removed. In practice, this served to remove long tails of very small clusters defined by the Leiden algorithm.

#### Signal correlation analyses

We converted footprint data files into a vector of footprint midpoint abundance for sequences S1 and S2 by summing footprint midpoint occurences and normalizing against the total number of footprints. We then correlated these vectors across replicate experiments using R for both correlation calculations and plotting associated scatterplots.

#### Trinucleosome analyses

Using processed footprint midpoint data files, we examined, for each footprinted fiber, the distances between all consecutive footprints sized between 100 and 200 bp, and plotted these distances against each other. All calculations were made on processed data tables generated using scripts described in the associated Jupyter notebook.

#### Autocorrelation analyses

Autocorrelations for *in vitro* and *in vivo* data were calculated using python, and then clustered as described above. All scripts for computing autocorrelation are available at the above link.

#### CTCF motif analyses

We examined the relative accessibility of 17 nucleotide windows tiling sequences S1 and S2 for each footprinted molecule before (native) and after (remodeled) remodeling, summarizing accessibility as a binary value thresholded on whether > 0.9 * the window length was accessible on a single molecule. We then stored these values in a data frame, and plotted the relative fractions of accessible windows against each other as log-odds values.

#### In vivo chromatin fiber analyses

All autocorrelation and clustering analyses were done as previously performed (Abdulhay et al., 2020). Autocorrelation and clustering were performed above. Nucleosome density enrichment plots were generated by estimating probability distributions for background (all molecules) and cluster-specific (clustered molecules) molecules, and computing log-odds from these distributions. All per-fiber nucleosome density measurements were calculated as above. Fisher’s Exact enrichment tests were carried out using scipy in Python as in Abudlhay et al (2020). All *p*-values calculated were then corrected using a Storey *q*-value correction, using the qvalue package in R (Storey and Tibshirani, 2003). Multiple hypothesis correction was performed for all domain-level Fisher’s tests (including ATAC peak analyses) and cutoffs were made at *q* < 0.05.

Molecules falling within ENCODE-defined epigenomic domains were extracted using scripts published in Abdulhay et al (2020).

#### ATAC data reanalysis

SNF2hKO and WT ATAC-seq data (Barisic et al., 2019) were downloaded, remapped to mm10 using bwa, converted to sorted, deduplicated BAM files, and then processed using macs2 to define accessibility peaks. Peaks were then filtered for reproducibility using the ENCODE IDR framework, and reproducible peaks were preserved for downstream analyses. Reproducible peaks for SNF2hKO and WT samples were pooled and merged using bedtools merge, and then used to generate count matrices using bedtools bamcoverage. Resulting count matrices for replicate experiments were then fed into DESeq2 to define statistically significant differentially accessible peaks with an adjusted *p-*value cutoff of 0.05.

#### In vivo TF binding analyses

Molecules containing previously-defined (Ramani et al., 2019) ENCODE-backed Ctcf bound motifs were extracted using scripts previously published (Abdulhay et al., 2020). Control molecules were obtained by randomly sampling a resource of Ctcf motif matches provided by Vierstra et al (2021). Single-molecule signals centered at the predicted motif center were stored as an array and clustered as above to obtain the clusters shown in the text. Enrichment analyses and associated multiple-hypothesis correction were performed as above for all enrichment tests performed for this array of Ctcf sites.

#### Satellite sequence analyses

Detecing mouse minor (centromeric) and major (pericentromeric) satellite is challenging because of the similarity of these two sequences (including internal / self-similarity). The latter is also an issue with the telomere repeat. To use BLAST to find matches to these sequences, the output must be processed to remove overlapping matches, which is done here heuristically using an implementation of the weighted interval scheduling dynamic programming algorithm that seeks to optimize the summed bitscores for non-overlapping matches to all three sequences (minor satellite, major satellite, and telomeres). This is not a perfect solution to the problem, in part because it treats the alignment for the three different repeats as effectively equivalent and we do not believe the alignments produced by BLAST are optimal compared to *e.g*. Smith-Waterman alignment, and the attendant fuzziness introduced may lead to removal of a small fraction of *bona fide* matches.

Given the similarity of major and minor satellite sequences in particular, using the DFAM minor (SYNREP_MM, accession DF0004122.1) and major (GSAT_MM, accession: DF0003028.1) satellite consensus sequences, which both exceed well-established monomer lengths of ~120 bp (minor) and ~234 bp (major), produces too many overlapping hits. Thus, we used more representative sequences from Genbank, specifically M32564.1 for major satellite, and X14462.1 for minor satellite. The telomere repeat sequence was constructed by pentamerizing the telomere repeat (*i.e*. [TTAGGG] x 5). All code used for these analyses is deposited at the above GitHub link.

## DATA AVAILABILITY

All processed data will be made available at Zenodo (https://doi.org/10.5281/zenodo.5770727). Raw data and a portion of the processed data will be uploaded to GEO at GEO accession XXXXX. All scripts and notebooks used for data analysis in this study will be made available at https://github.com/RamaniLab/SAMOSA-ChAAT.

## ACKNOWLEDGEMENTS

The authors thank Srinivas Ramachandran (CU Denver), Jay Sarthy (Fred Hutchinson Cancer Research Center), and Kliment Verba (UCSF) for comments on the manuscript. The authors thank Daniele Canzio (UCSF), Elphege Nora (UCSF), Hiten Madhani (UCSF), and the greater basic sciences faculty at UCSF for helpful discussions. This work was funded by grant DP2-HG012442 (to VR), grant U01-DK127421 (to GJN and VR), and grant R35-GM127020 (to GJN). Portions of this work were funded through the gracious support of the UCSF Program for Breakthrough Biomedical Research and the Sandler Fellows program.

## SUPPLEMENTARY FIGURES

**Supplementary Figure 1:**
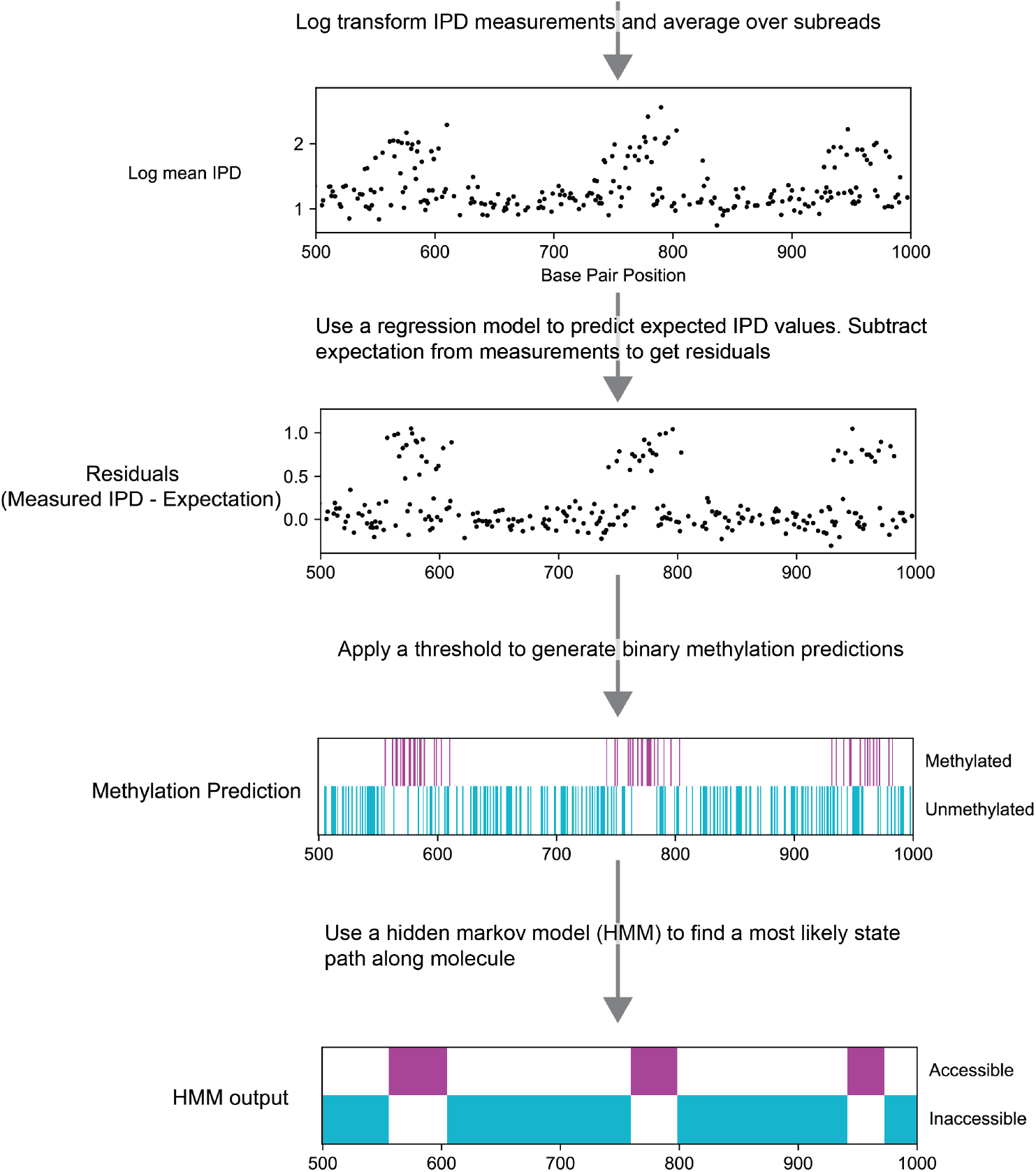
Computational pipeline for inferring DNA accessibility from measured inter-pulse distance (IPD). Shown is example data for a portion of a methylated molecule containing nucleosomes assembled onto regularly spaced Widom 601 sequences. The pipeline starts with log_10_ transforming the IPD measurements and averaging over all subreads. Next, to reduce noise from DNA sequence effects and inter-molecular variation, a neural network regression model that was trained on unmethylated DNA is used to regress out the expect IPD at each adenine. The regression model takes into account the DNA sequence context as well as molecule level IPD distribution measurements. The residuals show greater signal, and a threshold is then applied to the residuals to get binary methylation predictions. A hidden Markov model (HMM) is then used to synthesize the information from all adenines across the molecule into a single trace of accessible and inaccessible regions. The HMM model uses the frequency at which adenines in different sequence contexts were methylated in unmethylated and fully methylated control molecules to set expectations for observing methylation in accessible and inaccessible regions of chromatin. This HMM output was used for all downstream analyses.

**Supplementary Figure 2:**
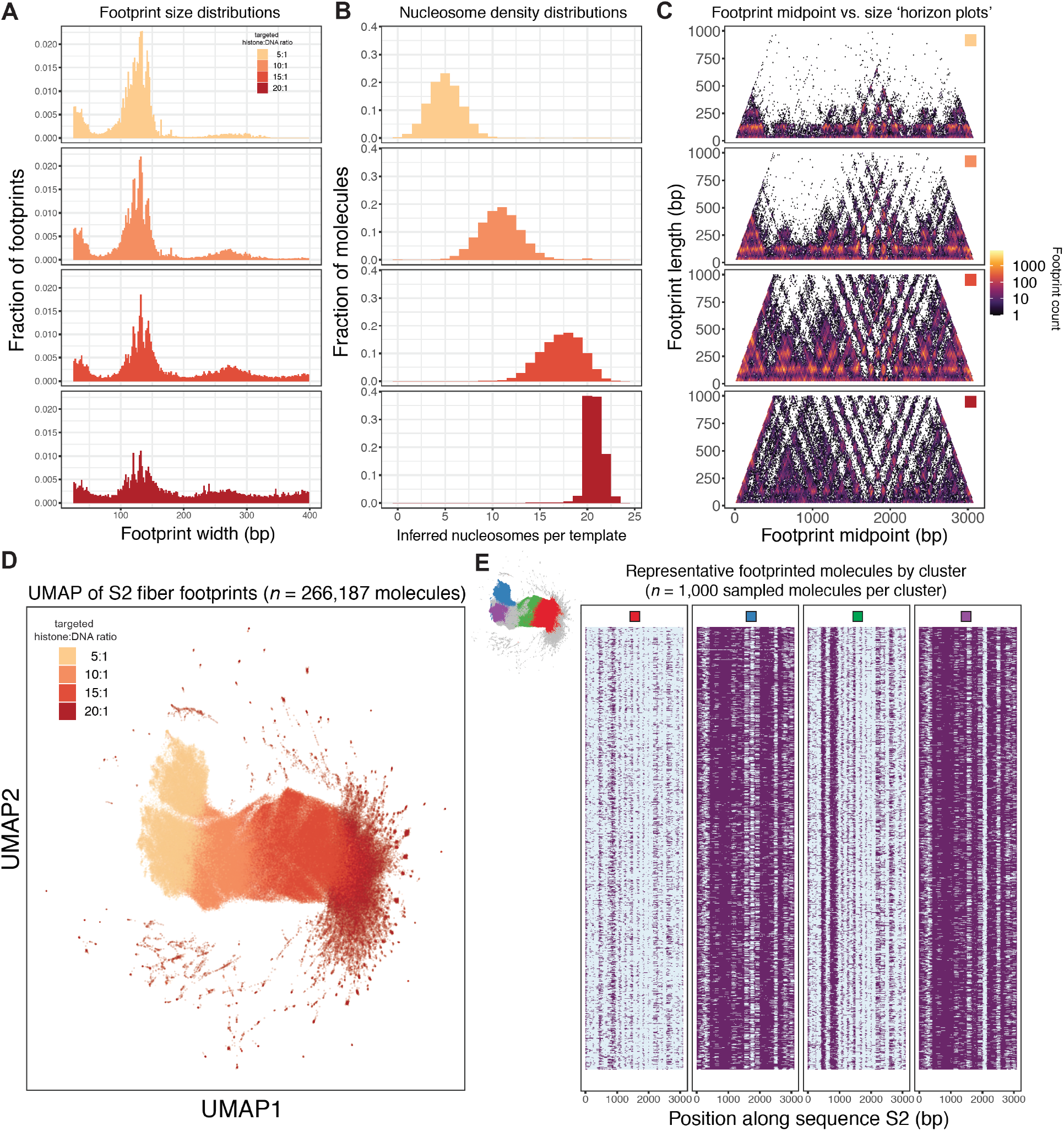
SAMOSA-ChAAT generalizes to other genomic sequences of interest. **A.)** As in **Figure 1B**, but for a completely different murine sequence (‘S2’). Footprint sizes from SAMOSA-ChAAT experiments carried out at varying nucleosome densities follow closely with expected nucleosome sizes, plus expected ‘breathing’ of DNA around the histone octamer, with the extent of breathing decreasing as nucleosome density increases. **B.)** SAMOSA-ChAAT data enables direct estimation of the absolute number of nucleosomes per footprinted S2 fiber. **C.)** Footprint length vs. midpoint ‘horizon’ plots for footprinted S2 fibers. **D.)** UMAP dimensionality reduction of S2 fiber accessibility data. **E.)** Visualization of a subset of sampled molecules following Leiden clustering of single molecule data.

**Supplementary Figure 3:**
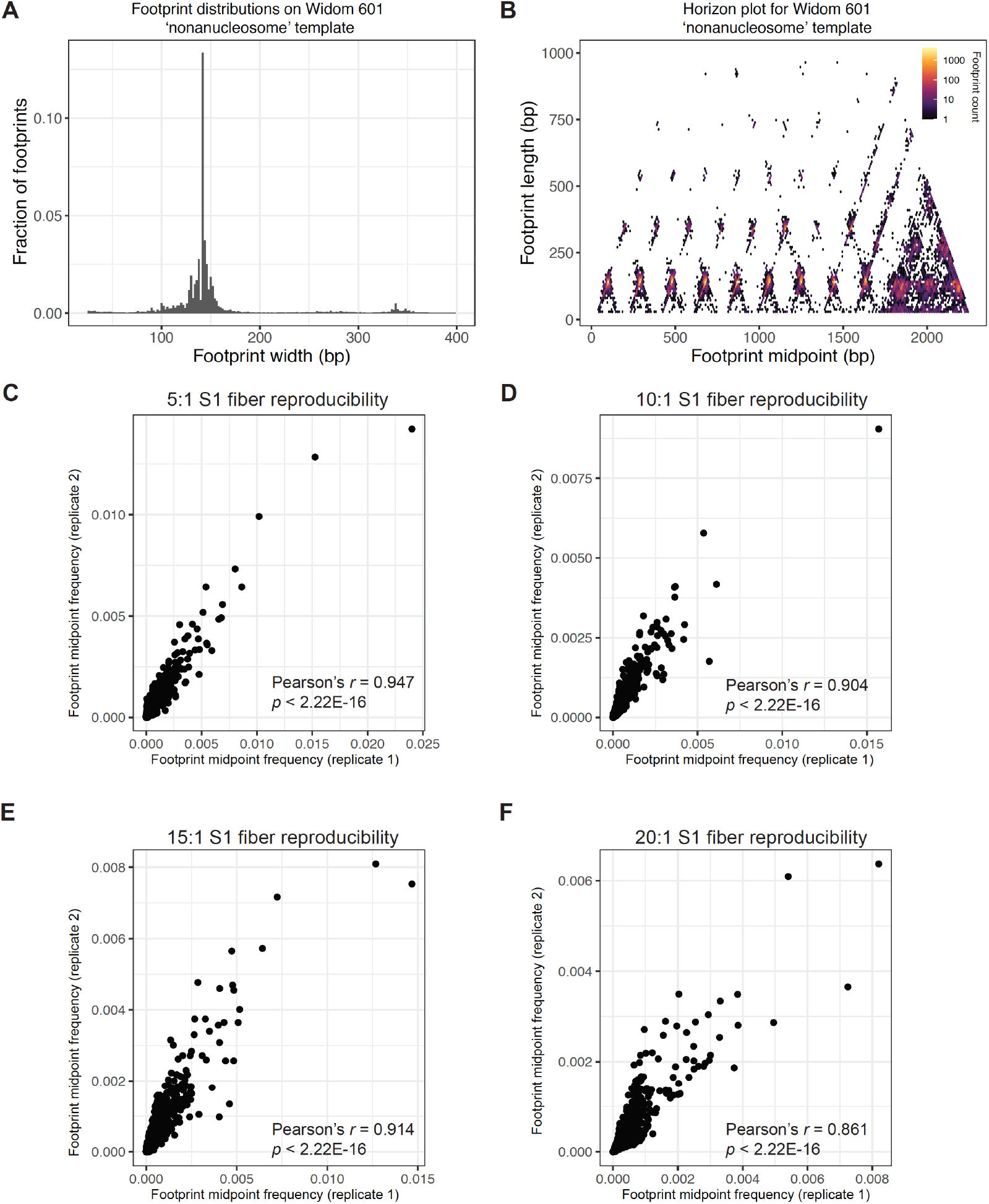
SAMOSA-ChAAT is accurate and reproducible. **A.-B)** Widom 601 nonanucleosomal fiber data from (Abdulhay et al., 2020) was reprocessed using the NN-HMM. Called footprints are the expected length of 601-assembled nucleosomes (**A**), and horizon plots reveal positioned nucleosomes at expected positions **(B). C-F.)** Correlation of footprint abundances for S1 fibers of each density across two replicates (different remodeling reactions on fibers generated from different salt gradient dialysis preps). SAMOSA-ChAAT measurements are highly reproducible.

**Supplementary Figure 4.**
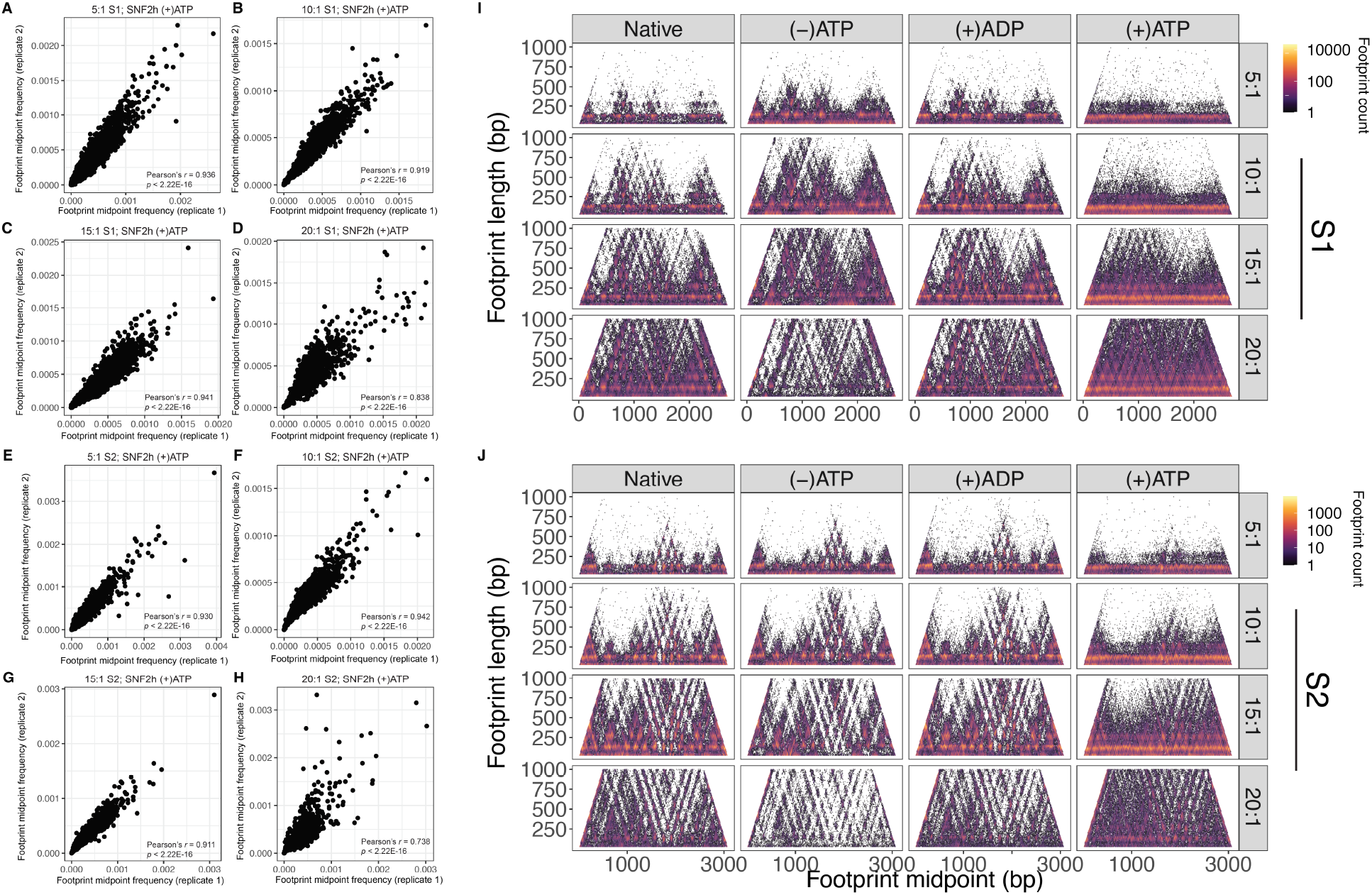
Reproducibility of SAMOSA-ChAAT remodeling experiments and horizon plots for all catalytic conditions tested. **A-H.)** Scatter plots and associated Pearson’s *r* values for correlations between two biological replicate remodeling experiments, for each density tested, for both S1 (**A-D**) and S2 (**E-H**) fibers. **I-J.**) Horizon plots for S1 (**I**) and S2 (**J**) fibers, for native, pre-catalytic, (+)ADP, and remodeled fibers (all averages are over single-turnover experiments; multi-turnover data is omitted for this visualization).

**Supplementary Figure 5:**
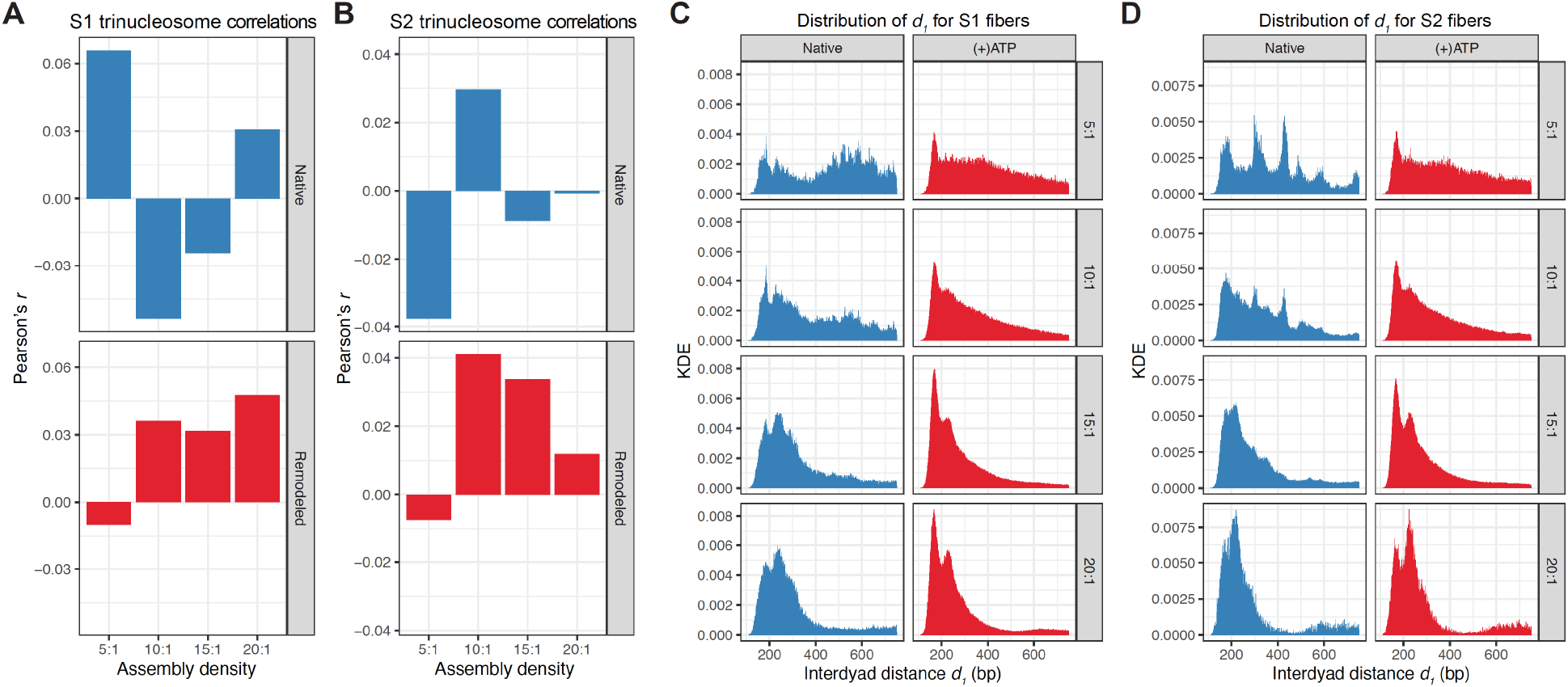
Nucleosome spacing correlation is slightly impacted by chromatin remodeling, and dinucleosome distances are not fixed by SNF2h remodeling. **A.-B.)** Bar chart representation of Pearson’s *r* values for each assembly density, for unremodeled (blue) and remodeled (red) S1 (**A**) and S2 (**B**) fibers. Correlations are globally very low (indicating little coupling between the two distances), and is slightly impacted in both directions upon remodeling, though the effects are small. **C-D.)** Histograms (binsize = 1 bp) for unremodeled (blue) and remodeled (red) interdyad distances (*i.e. d*_1_) for S1 (**C**) and S2 (**D**) fibers.

**Supplementary Figure 6:**
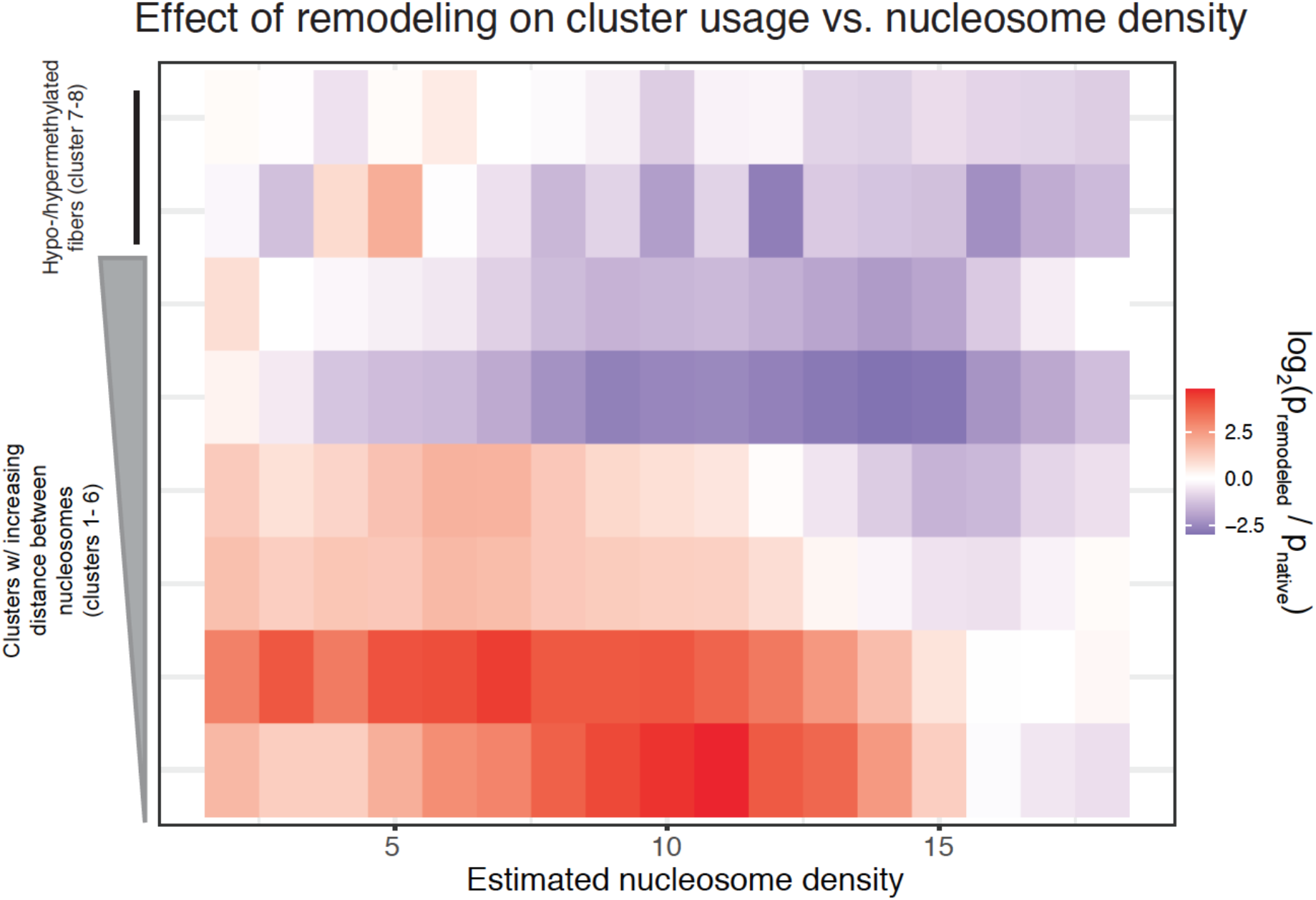
SNF2h remodeling alters the probability of observing specific chromatin fiber structures, and remodeling outcomes are highly heterogeneous. Log-odds visualization capturing the likelihood that chromatin fiber states are increased (red) or decreased (blue) in representation with respect to native, unremodeled fibers. p_remodeled_: fraction of molecules of a given cluster for each nucleosome density estimate after SNF2h remodeling; p_native_: fraction of molecules of a given cluster for each nucleosome density estimate before SNF2h remodeling.

**Supplementary Figure 7.**
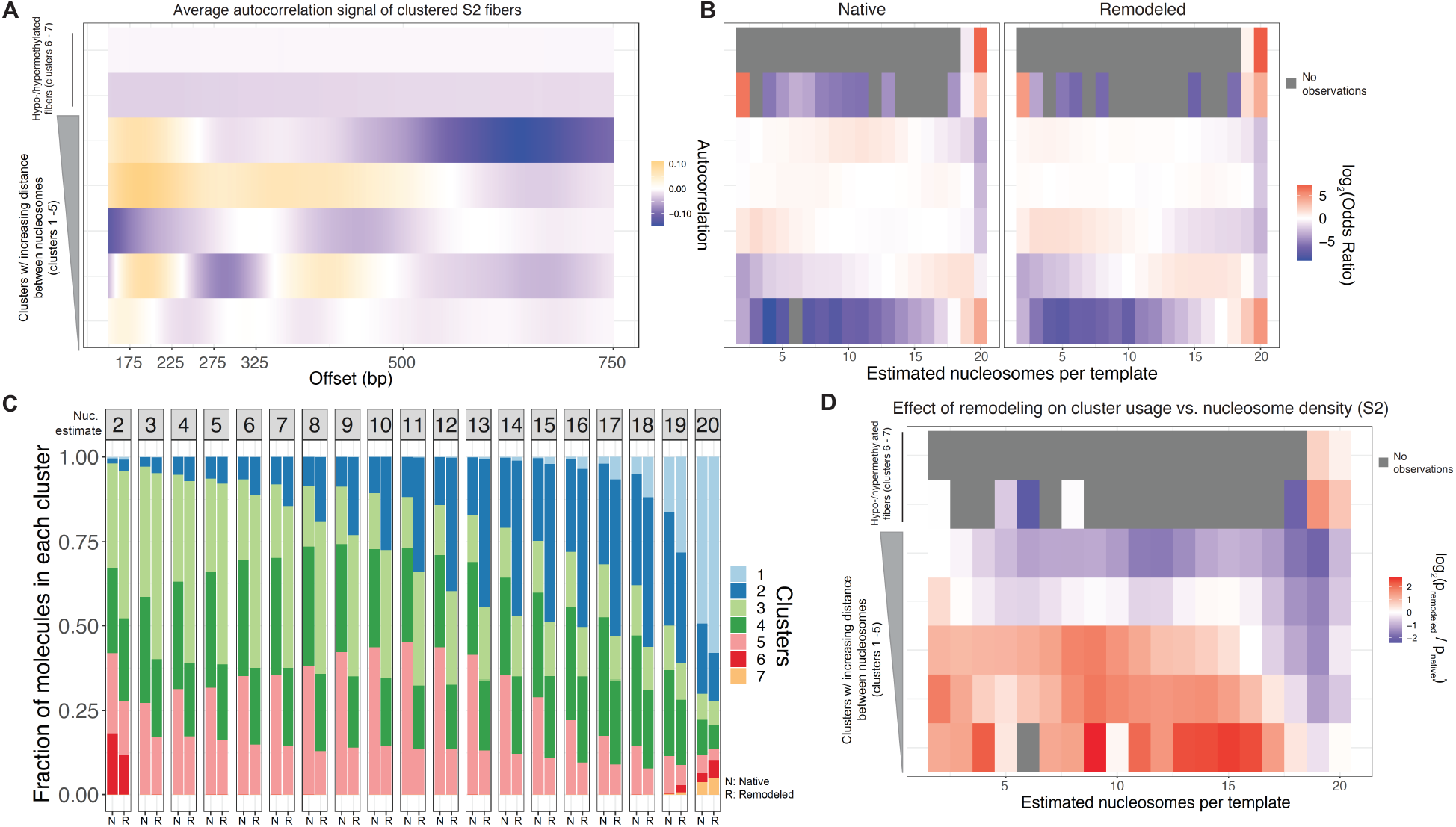
Autocorrelation clustering analysis for fiber S2. **A.)** As in **Figure 4B**, but for fiber S2; unsupervised Leiden clustering of single-molecule autocorrelations yields 7 clusters. **B.)** As in **Figure 4C**, Fishers exact effect sizes for cluster enrichment as a function of estimated chromatin density. **C.)** As in **Figure 4D**, a stacked bar-chart representation of the relative abundance of each fiber-type, stratified by estimated chromatin fiber density, for S2 fibers. **D.)** Log-odds visualization capturing the likelihood that chromatin fiber states are increased (red) or decreased (blue) in representation with respect to native, unremodeled S2 fibers. p_remodeled_: fraction of molecules of a given cluster for each nucleosome density estimate after SNF2h remodeling; p_native_: fraction of molecules of a given cluster for each nucleosome density estimate before SNF2h remodeling.

**Supplementary Figure 8:**
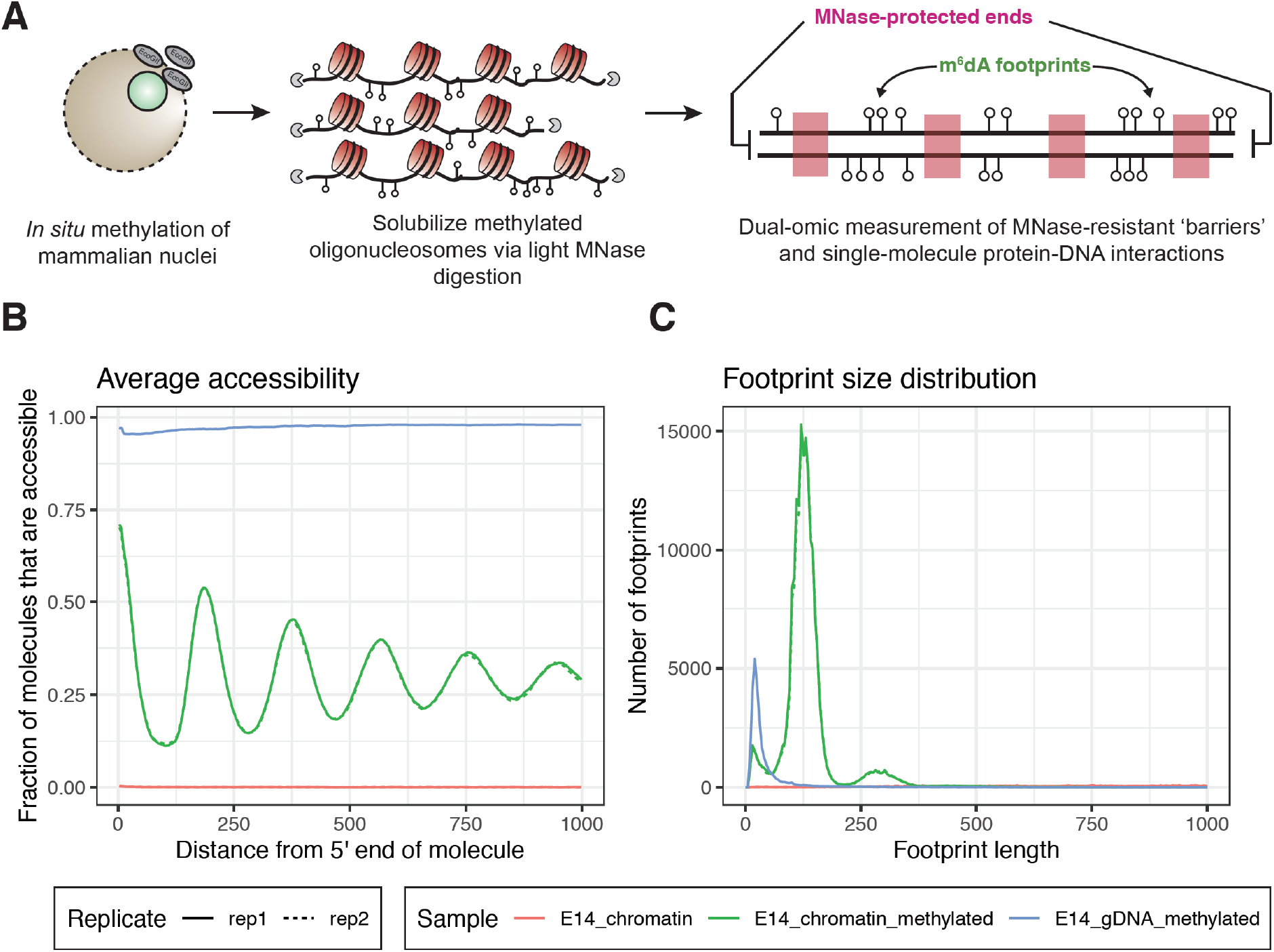
An improved, *in situ* SAMOSA assay for profiling single-fiber chromatin structure *in vivo*. **A.)** We improved on our previously published SAMOSA protocol by performing EcoGII methylation in intact nuclei, which we then digest with a limited MNase digestion to liberate oligonucleosomes. These molecules are then sequenced on the PacBio Sequel II platform and harbor two information types: MNase cuts that mark the position of ‘barriers’ along the genome, and m^6^dA footprints that capture protein-DNA interactions. **B.)** Our NN-HMM can be applied to estimate chromatin accessibility on individual molecules. Shown here is data from E14 mESCs. Nucleosome periodicity is seen in footprinted chromatin, but not in positive (methylated naked DNA) and negative (unmethylated E14 gDNA) controls. The 5’ and 3’ ends of molecules are massively enriched for MNase-defined ‘barriers’ (generally, the edge of nucleosome core particles). **C.)** The NN-HMM can predict footprint sizes, which range from nucleosome length, to subnucleosomal protections indicative of transcription factor-DNA interactions.

**Supplementary Figure 9:**
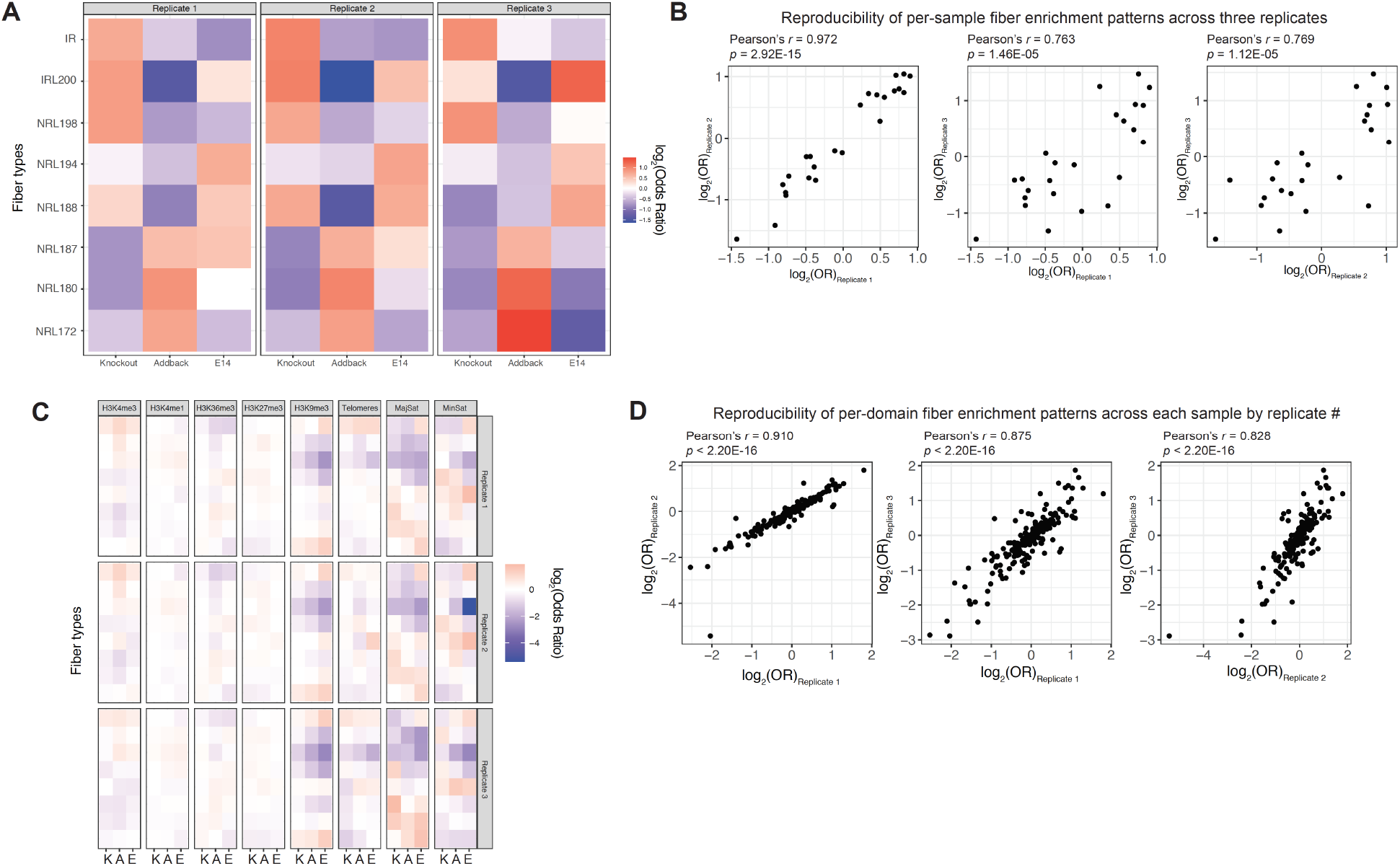
Reproducibility of *in vivo* SAMOSA data from knockout, addback, and E14 mESCs. **A.)** Fisher’s exact test results for sample-level fiber enrichment stratified by biological replicate. **B.)** Pearson’s *r* correlations, *p*-values, and associated scatter plots of effect sizes for three biological replicates. **C.)** Fisher’s exact test results for domain-level fiber enrichment stratified by biological replicate. **D.)** Pearson’s *r* correlations, *p*-values, and associated scatter plots of effect sizes for domain-level analyses, across three biological replicates.

**Supplementary Figure 10:**
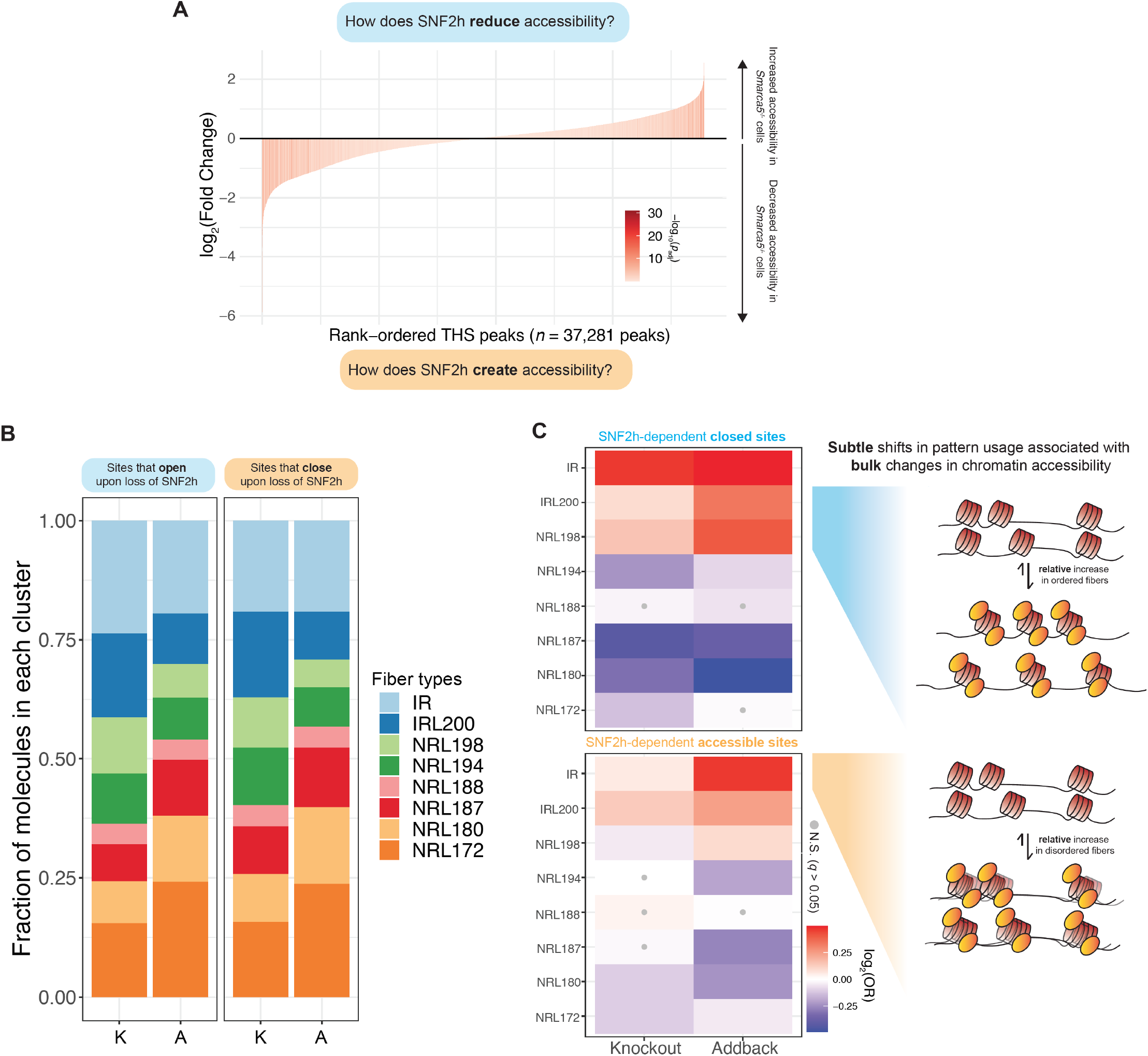
Relative abundance of each chromatin fiber type at differentially accessible ATAC-seq peaks. **A.)** We re-analyzed published ATAC-seq data to determine statistically significant Tn5-hypersensitive sites (THSs) that are genetically dependent on SNF2h for maintaining chromatin accessibility patterns. Regions that open upon SNF2h-loss can be used to study how SNF2h reduces bulk chromatin accessibility, while regions that close upon SNF2h-loss can be used to study how SNF2h maintains bulk chromatin accessibility. **B.)** A stacked bar chart representation of the relative usage of each fiber type in (left) regions that are significantly more accessible in knockout cells versus wildtype cells, and (right) regions that are significantly less accessible in knockout cells versus wildtype cells. **C.)** Fisher’s exact test results for fibers falling in regions that open (top) and regions that close (bottom) in knockout cells by ATAC-seq. Upon reintroduction of SNF2h in addback cells, regions that open increase representation of evenly-spaced chromatin fibers, while regions that close increase the relative representation of irregular fibers.

**Supplementary Figure 11:**
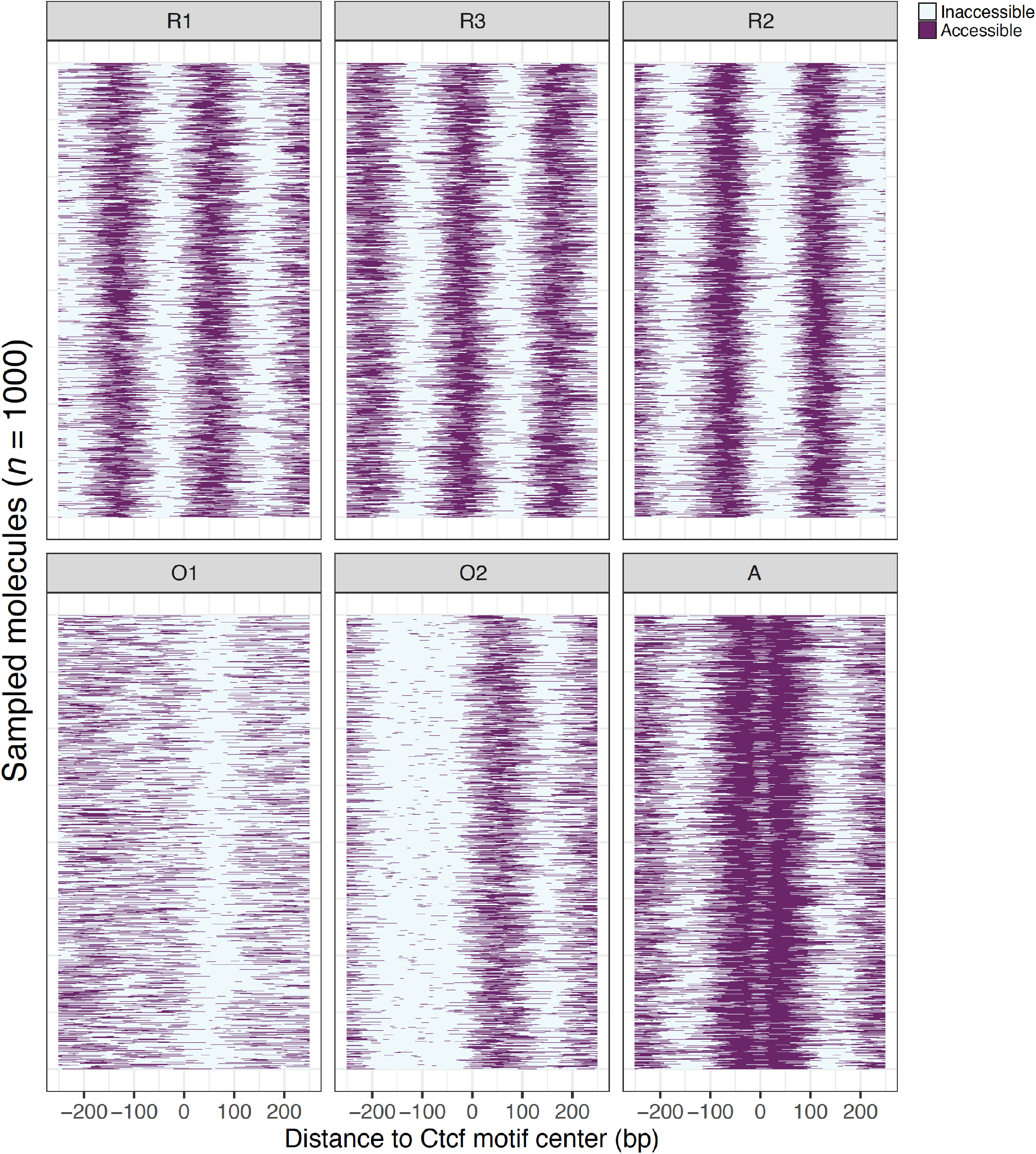
Single-fiber nucleosome occupancy patterns around predicted Ctcf binding sites *in vivo*. 1000 randomly sampled chromatin fibers, with accessibility signal centered around the predicted Ctcf binding motif, for each of six different nucleosome occupancy patterns obtained through unsupervised Leiden clustering. R1 – R3 are regularly phased nucleosomes, O1 – O2 are ‘irregular’ nucleosome occupancy patterns that appear to occlude the cognate binding motif for Ctcf, and A represents an ‘accessible’ state. A fraction of A molecules display a ‘footprint’ of unmethylated DNA precisely over the Ctcf binding site, indicative of a molecule where Ctcf was bound during the footprinting reaction.

**Supplementary Figure 12:**
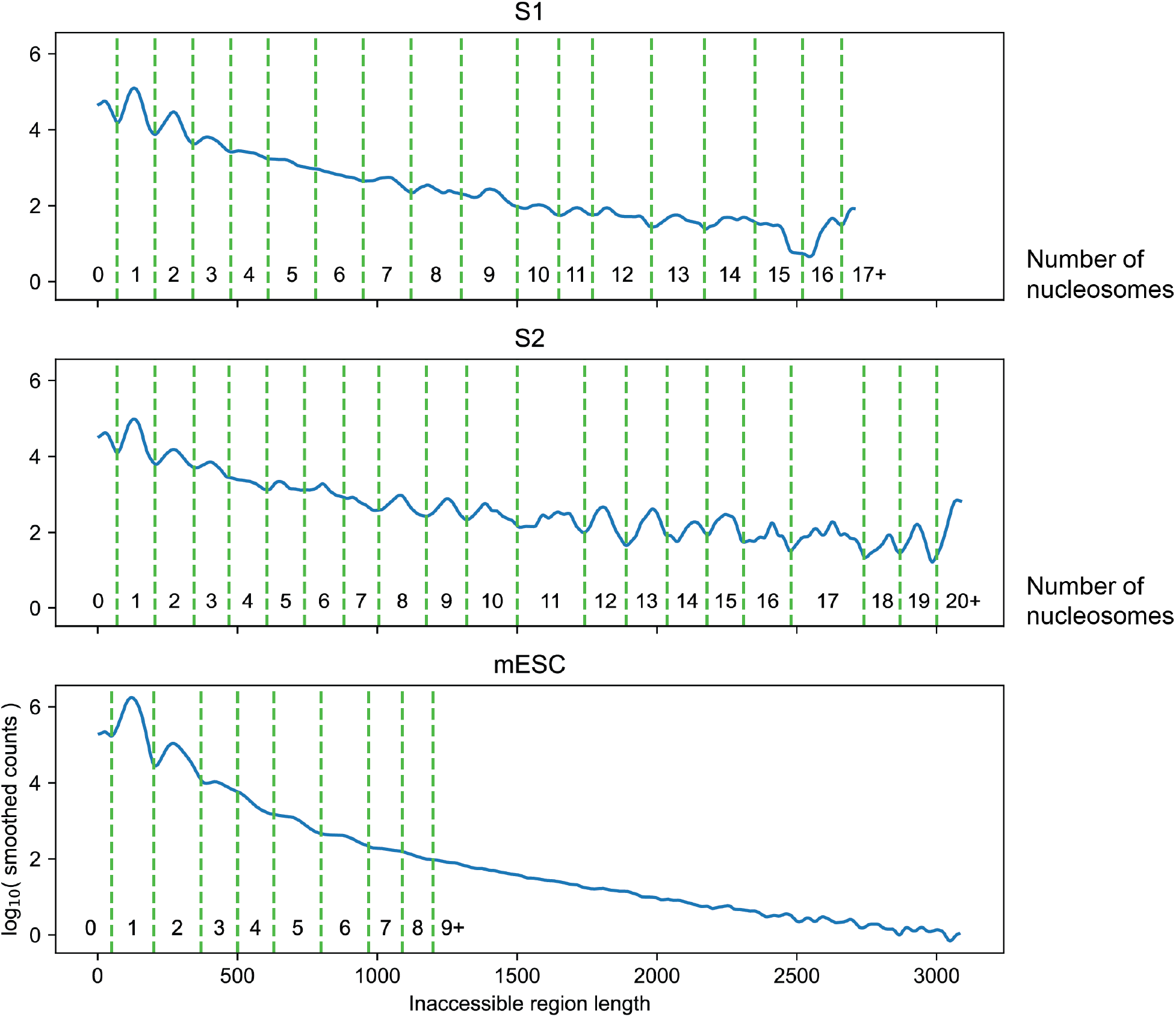
Cutoffs for counting nucleosomes in inaccessible regions based on length. To estimate the number of nucleosomes on each DNA molecule, cutoffs were defined to delineate between the number of estimated nucleosomes within an inaccessible region. Green dashed lines show the cutoffs, and the numbers below indicate the number of nucleosomes that sized region is counted as. Different cutoffs were used for S1, S2, and mESC molecules, based on the distributions and peaks in region length for each.

## SUPPLEMENTARY TABLES

**Supplementary Table 1:**
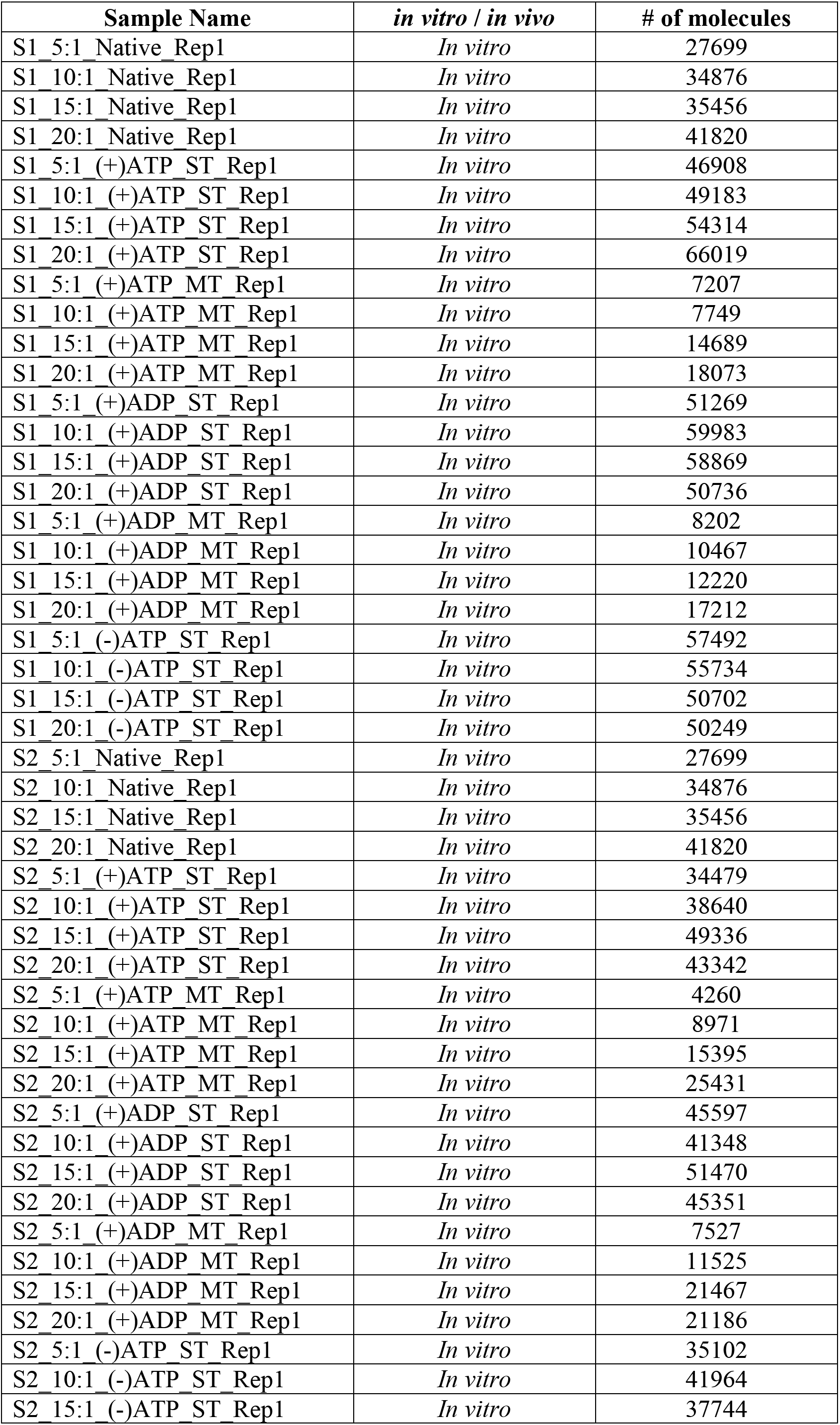
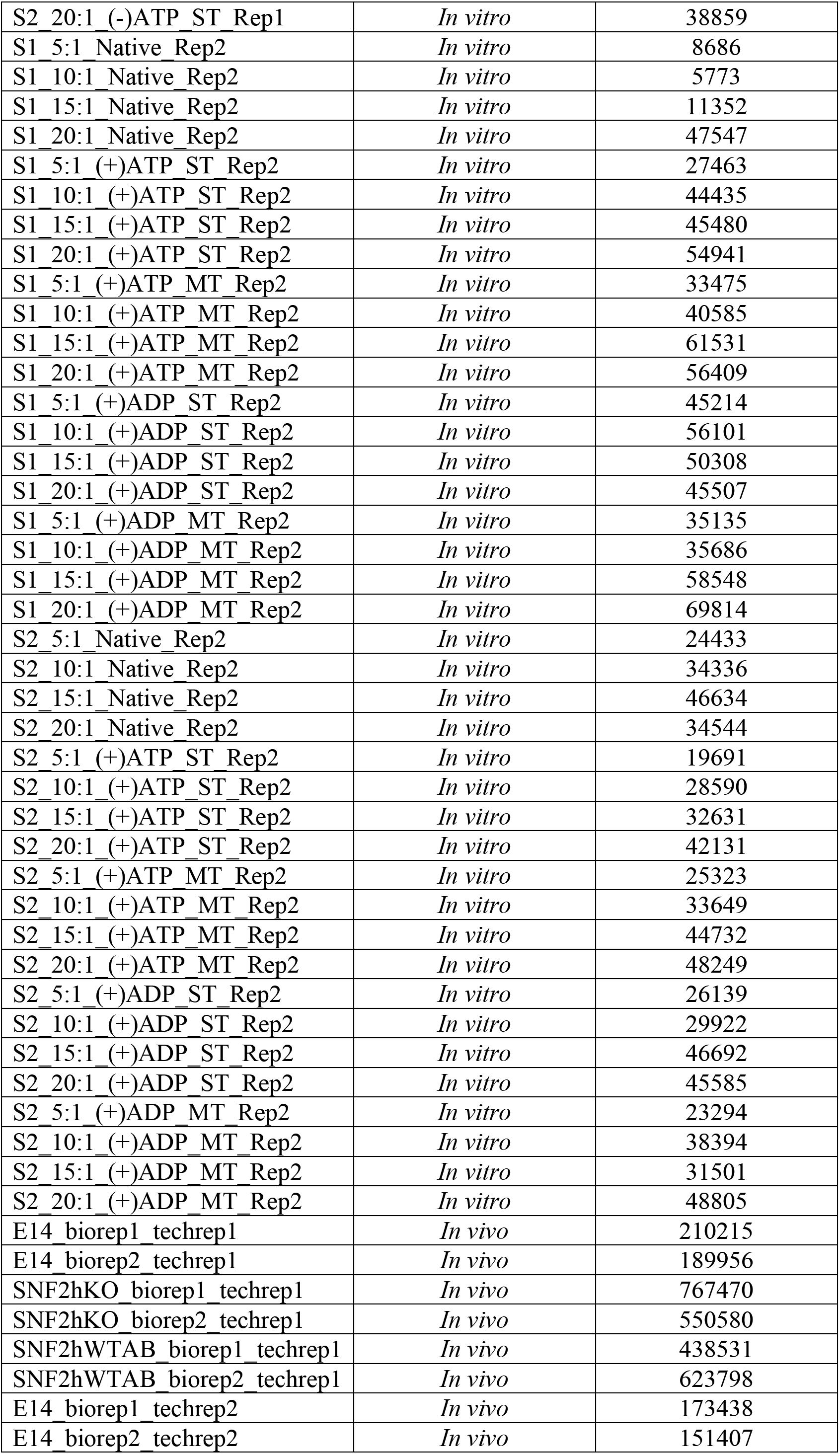
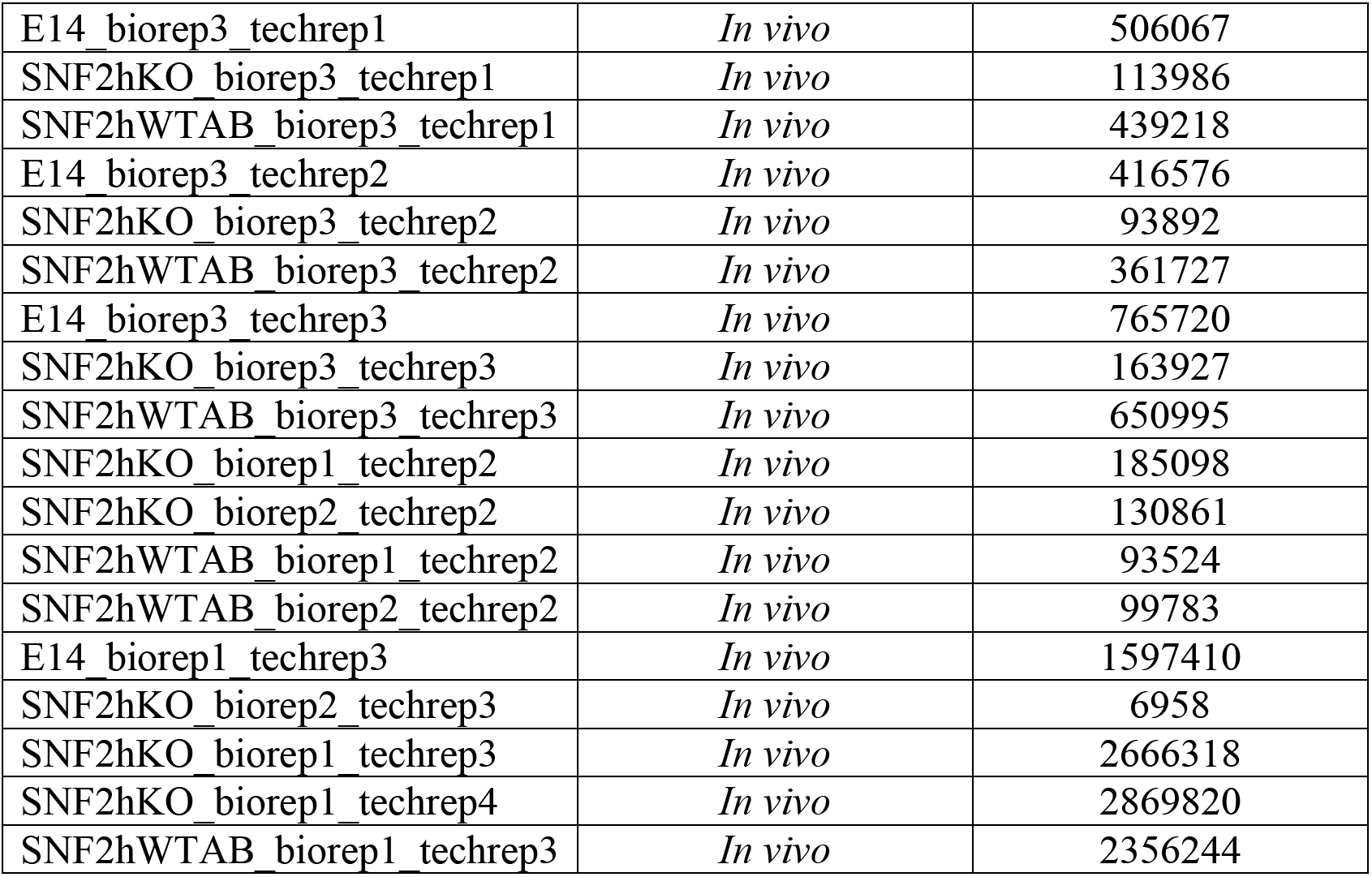
Summary of sequencing depths for all experiments performed in this study. For *in vivo* samples, ‘biorep’ refers to different biological replicate experiments, while ‘techrep’ refers to ‘technical replicates’ where the same sequencing library was sequenced on multiple PacBio Sequel II runs.

**Supplementary Table 2:** Summary of average nucleosomes and standard deviation for all *in vitro* experiments. ST refers to single-turnover remodeling reaction conditions, while MT refers to multi-turnover reaction conditions. Data are shown as mean ± standard deviation.

| Fiber Type | Density | Nucleosomes / template | Remodeling status |
| --- | --- | --- | --- |
| S1 | 5:1 | 4.00 ± 1.80 | Native |
| S1 | 10:1 | 7.93 ± 2.22 | Native |
| S1 | 15:1 | 12.6 ± 2.16 | Native |
| S1 | 20:1 | 15.8 ± 1.60 | Native |
| S2 | 5:1 | 5.17 ± 2.00 | Native |
| S2 | 10:1 | 11.0 ± 2.16 | Native |
| S2 | 15:1 | 17.3 ± 2.19 | Native |
| S2 | 20:1 | 20.8 ± 0.918 | Native |
| S1 | 5:1 | 3.77 ± 1.93 | (+) ATP (ST) |
| S1 | 10:1 | 7.28 ± 2.04 | (+) ATP (ST) |
| S1 | 15:1 | 11.8 ± 2.01 | (+) ATP (ST) |
| S1 | 20:1 | 15.5 ± 1.80 | (+) ATP (ST) |
| S2 | 5:1 | 4.68 ± 1.93 | (+) ATP (ST) |
| S2 | 10:1 | 9.92 ± 2.11 | (+) ATP (ST) |
| S2 | 15:1 | 15.6 ± 2.10 | (+) ATP (ST) |
| S2 | 20:1 | 20.9 ± 0.987 | (+) ATP (ST) |
| S1 | 5:1 | 4.62 ± 1.88 | (-) ATP (ST) |
| S1 | 10:1 | 10.2 ± 2.22 | (-) ATP (ST) |
| S1 | 15:1 | 15.4 ± 1.65 | (-) ATP (ST) |
| S1 | 20:1 | 17.0 ± 0.918 | (-) ATP (ST) |
| S2 | 5:1 | 5.22 ± 2.00 | (-) ATP (ST) |
| S2 | 10:1 | 11.4 ± 2.10 | (-) ATP (ST) |
| S2 | 15:1 | 18.5 ± 1.72 | (-) ATP (ST) |
| S2 | 20:1 | 20.8 ± 0.909 | (-) ATP (ST) |
| S1 | 5:1 | 3.60 ± 1.45 | (+) ATP (MT) |
| S1 | 10:1 | 6.96 ± 1.88 | (+) ATP (MT) |
| S1 | 15:1 | 11.9 ± 1.87 | (+) ATP (MT) |
| S1 | 20:1 | 15.2 ± 1.50 | (+) ATP (MT) |
| S2 | 5:1 | 5.37 ± 1.72 | (+) ATP (MT) |
| S2 | 10:1 | 11.2 ± 1.84 | (+) ATP (MT) |
| S2 | 15:1 | 18.0 ± 1.81 | (+) ATP (MT) |
| S2 | 20:1 | 20.7 ± 0.818 | (+) ATP (MT) |
| S1 | 5:1 | 4.07 ± 1.78 | (+) ADP (ST) |
| S1 | 10:1 | 8.02 ± 1.90 | (+) ADP (ST) |
| S1 | 15:1 | 12.8 ± 1.81 | (+) ADP (ST) |
| S1 | 20:1 | 16.1 ± 1.39 | (+) ADP (ST) |
| S2 | 5:1 | 5.39 ± 2.13 | (+) ADP (ST) |
| S2 | 10:1 | 11.3 ± 2.22 | (+) ADP (ST) |
| S2 | 15:1 | 17.4 ± 1.99 | (+) ADP (ST) |
| S2 | 20:1 | 20.8 ± 0.965 | (+) ADP (ST) |
| S1 | 5:1 | 3.66 ± 1.65 | (+) ADP (MT) |
| S1 | 10:1 | 6.91 ± 1.77 | (+) ADP (MT) |
| S1 | 15:1 | 11.9 ± 1.88 | (+) ADP (MT) |
| S1 | 20:1 | 15.3 ± 1.47 | (+) ADP (MT) |
| S2 | 5:1 | 5.46 ± 2.01 | (+) ADP (MT) |
| S2 | 10:1 | 11.3 ± 1.84 | (+) ADP (MT) |
| S2 | 15:1 | 18.2 ± 1.79 | (+) ADP (MT) |
| S2 | 20:1 | 20.7 ± 0.797 | (+) ADP (MT) |

